# Rescue of pATOM36-depleted T. brucei by human MTCH1/2 reveals common features of protein insertases

**DOI:** 10.64898/2026.02.13.705746

**Authors:** Stephan Berger, Christoph von Ballmoos, André Schneider

## Abstract

Alpha-helically anchored proteins of the mitochondrial outer membrane depend on dedicated insertases, including the MIM complex in yeast, pATOM36 in trypanosomes, and MTCH1/MTCH2 in humans. These insertases lack sequence homology and arose by convergent evolution. Here we show that MTCH1 and an N-terminally modified MTCH2 can replace the OMM biogenesis function of trypanosomal pATOM36, demonstrating functional compatibility across a large phylogenetic distance. Moreover, like other insertases, pATOM36 has lipid scramblase activity and likely induces local membrane thinning. Structural modeling reveals that even though they have inverted membrane topologies, pATOM36 and MTCH1/MTCH2 adopt a similar architecture. They form a hydrophilic cavity comprising five transmembrane helices that is open to both the cytosol and laterally to the membrane core. The same features also extend to the large phylogenetically unrelated YidC-like insertase family of bacterial origin. Together, these findings suggest that membrane protein insertion is constrained to a small number of viable mechanistic solutions, providing evidence for “deterministic” evolution acting at the molecular level.

## Introduction

All mitochondria have an own genome, which reflects their bacterial ancestry. However, this genome encodes only few proteins most of which are involved in oxidative phosphorylation. Thus, more than 95% of the 1000-1500 mitochondrial proteins are encoded in the nucleus, synthesized in the cytosol, and subsequently imported into the organelle^1–3^. Irrespective of their intracellular localization essentially all imported mitochondrial proteins use the protein translocase of the outer membrane (TOM) as their entry gate into the organelle. This includes integral outer mitochondrial membrane (OMM) β-barrel proteins which are first translocated across the OMM by the TOM complex and subsequently inserted from the intermembrane space (IMS) side into the OMM by the sorting and assembly machinery (SAM)^4,5^. The OMM also contains approximately 100 proteins that are α-helically anchored. Most of these proteins have a single transmembrane domain (TMD), although a few multipass TMD-containing OMM proteins also exist^6–8^.

In the yeast *Saccharomyces cerevisiae,* most α-helically anchored OMM proteins are inserted by a dedicated OMM insertase termed mitochondrial import machinery (MIM) complex. The MIM complex consists of the two C-terminally anchored OMM proteins, Mim1 (13 kDa) and Mim2 (11 kDa)^9–11^. It can either directly insert its substrate into the OMM or in some cases may co-operate with the import receptors Tom70 and Tom20 of the TOM complex. While the exact structure and stoichiometry of the MIM complex is unknown, it appears to contain more copies of Mim1 than of Mim2^11–13^. In addition, the MIM complex is important for TOM complex assembly because it is required for OMM insertion of α-helically anchored TOM subunits (Fig. 1A)^9,12^.

**Figure 1:**
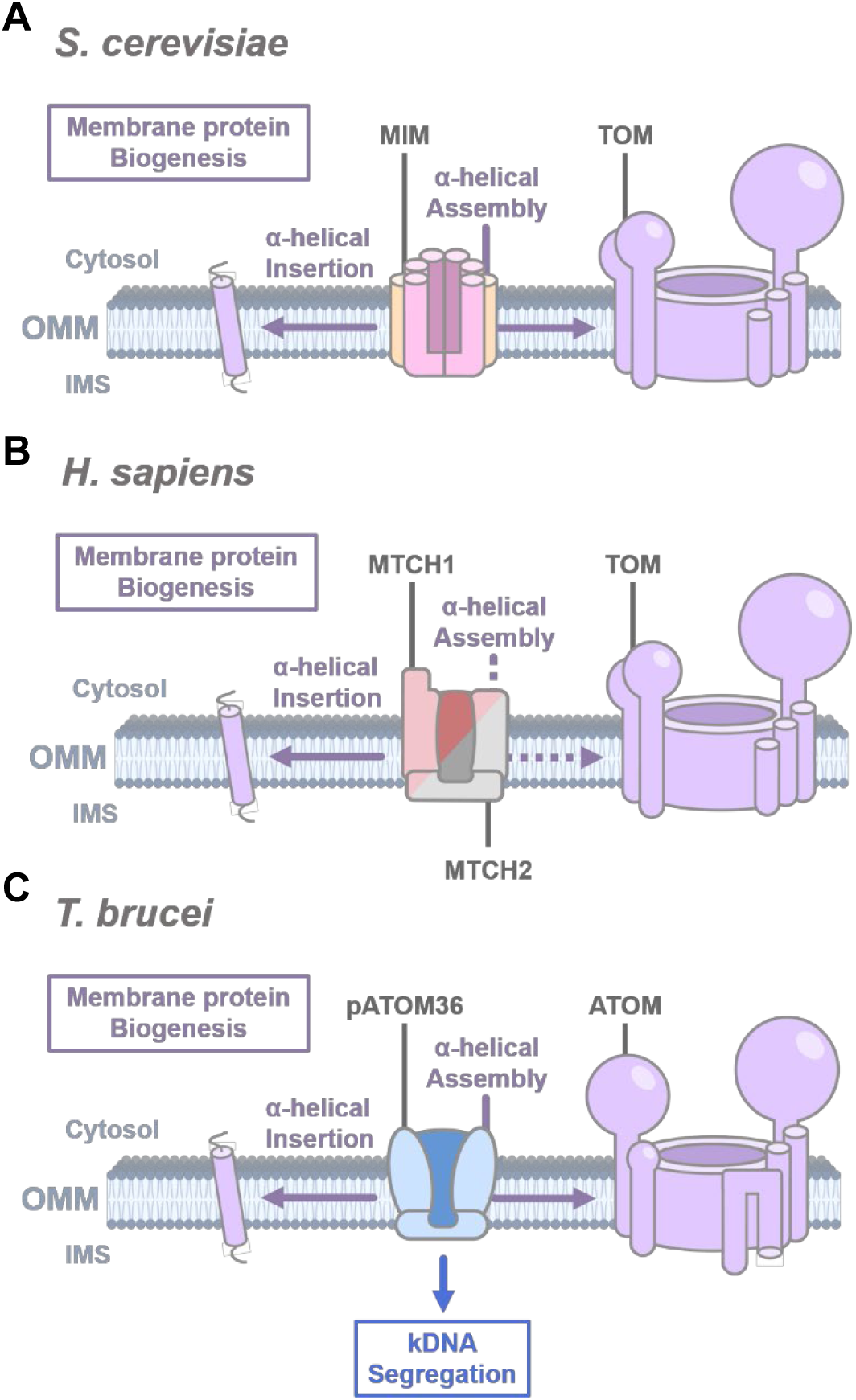
Membrane protein biogenesis is regulated by distinct molecular machineries across different species. **A)** In the OMM of *S. cerevisiae*, the MIM complex, consisting of Mim1 (pink) and Mim2 (orange), mediates the insertion of α-helical membrane proteins and supports the assembly of the TOM complex (lavender). **B)** In *H. sapiens*, two independent insertases, MTCH1 (red) and MTCH2 (grey), mediate the integration of α-helical membrane proteins, although their involvement in TOM complex (lavender) assembly has not yet been established (dashed arrow). **C)** In *T. brucei* pATOM36 (light blue) functions as an insertase for α-helical membrane proteins and contributes to the assembly of the ATOM complex (lavender). In addition, pATOM36 has a TAC specific function and is involved in kDNA segregation (blue).

The TOM complexes of mammals and yeast, both of which belong to the opisthokonts, are highly conserved. However, Mim1 and Mim2 are only found in fungi^14^. In humans, most α-helically anchored integral OMM proteins, including subunits of the TOM complex, are inserted by the mitochondrial carrier homologue 2 (MTCH2) (33 kDa), a SLC25 family member^15^. MTCH1 (42 kDa) is the paralogue of MTCH2 and likely also has insertase activity^16^ (Fig. 1B). Previous studies suggested that MTCH1 and/or MTCH2 may also function in other processes such as apoptosis, ferroptosis, tumor progression, lipid metabolism, and mitochondrial fusion^17–20^. It is possible that these phenotypes are ultimately also connected to the insertase activity of MTCH1 and MTCH2.

The parasitic protozoan *Trypanosoma brucei* belongs to kinetoplastids within the Excavata supergroup and is essentially unrelated to opisthokonts. It has a highly diverged TOM complex, termed atypical TOM (ATOM)^14,21,22^. The ATOM complex has seven subunits: the pore-forming β-barrel membrane protein ATOM40 and six α-helically membrane-anchored proteins ATOM11, ATOM12, ATOM14, ATOM19, ATOM46, and ATOM69. Among these, ATOM46 and ATOM69 function as protein import receptors. ATOM40 and ATOM14 are orthologues of the opisthokont Tom40 and Tom22, respectively, whereas the remaining ATOM subunits are unique to kinetoplastids^22,23^.

Strikingly, trypanosomes lack orthologues of both Mim1/Mim2 and MTCH1/MTCH2^22–24^. Instead, it is the OMM protein peripheral ATOM36 (pATOM36) (36 kDa) that mediates the biogenesis of α-helically anchored OMM proteins, including many ATOM subunits^25,26^. Depletion of pATOM36 prevents assembly of the ATOM complex. Intriguingly, pATOM36 has a dual function^26,27^. It is also an essential subunit of the trypanosomal tripartite attachment complex (TAC) that physically links the single-unit mitochondrial genome of *T. brucei*, termed kinetoplast DNA (kDNA), with the basal body of the flagellum. The TAC mediates the segregation of the replicated kDNA during cytokinesis. Both the OMM protein biogenesis and the TAC function of pATOM36 are essential for normal growth of *T. brucei* (Fig. 1C)^26–29^. It appears that the MIM complex, MTCH2, and possibly MTCH1, as well as pATOM36, have the very same function in their respective systems: to insert a subset of α-helical membrane proteins, including (A)TOM subunits, into the OMM. For Mim1/Mim2, this was experimentally confirmed. Expression of pATOM36 in a yeast Mim1/Mim2 deletion strain restored both TOM complex assembly and wild-type growth^30^. Conversely, expression of Mim1/Mim2 in a pATOM36-depleted *T. brucei* cell line restored ATOM complex assembly.

However, growth was only partially restored, because Mim1/Mim2 cannot replace the TAC function of pATOM36^30^. It had also been shown that expression of MTCH1, but not of MTCH2, restored TOM complex assembly and wild-type growth in the Mim1/Mim2 yeast deletion strain^16^. These results are surprising, because the three insertases do not share any sequence homology. It suggests that the OMM insertases in the three systems arose independently by convergent evolution.

In the present study, we have used *in vivo* complementation to test whether mammalian MTCH1 or MTCH2 can replace the OMM protein biogenesis function of pATOM36 in *T. brucei*. Our results show this is indeed possible, provided that MTCH1 and MTCH2 are efficiently localized to the trypanosomal mitochondrion. Moreover, we show that recombinantly expressed and purified pATOM36 as many other protein insertases has lipid scramblase activity *in vitro*. Finally, we discuss the convergently evolved structural features that pATOM36 and MTCH1/MTCH2 share with other insertases.

## Results

### pATOM36-HA expression restores all deficiencies caused by pATOM36 depletion

pATOM36 is essential in both the procyclic and the bloodstream form life-cycle stages of *T. brucei* ^25,26,30^. In this work, we have used the previously established procyclic pATOM36 RNAi cell line that targets the 3’-UTR of the pATOM36 mRNA (pATOM36-3’UTR RNAi) ^26^. This cell line allows tetracycline-inducible depletion of endogenous pATOM36 together with simultaneous ectopic expression of a transgenic variant of the protein, provided that its mRNA carries a different 3ʹ UTR. The pATOM36-3ʹUTR RNAi cell line showed a growth arrest after three days of tetracycline addition (Fig. 2A). Moreover, Blue Native polyacrylamide gel electrophoresis (BN-PAGE) analysis indicated that pATOM36 depletion disrupted ATOM complex assembly. Whereas ATOM40-containing complexes range between 400 to 700 kDa in uninduced cells, they essentially disappeared after depletion of pATOM36. This phenotype is already clearly visible after two days and still increases after three days of RNAi induction (Fig. 2B, upper panels). Concomitant with ATOM complex depletion, the steady-state levels of the previously identified pATOM36 substrates ATOM46, ATOM19, and ATOM14 are reduced, as assessed by denaturing SDS-PAGE. The levels of ATOM40, however, which is not a substrate of pATOM36, remained unchanged (Fig. 2B, lower panels). ATOM69 showed intermediate behavior and is only slightly depleted. In summary, these results are in complete agreement with previously reported findings^26^.

**Figure 2:**
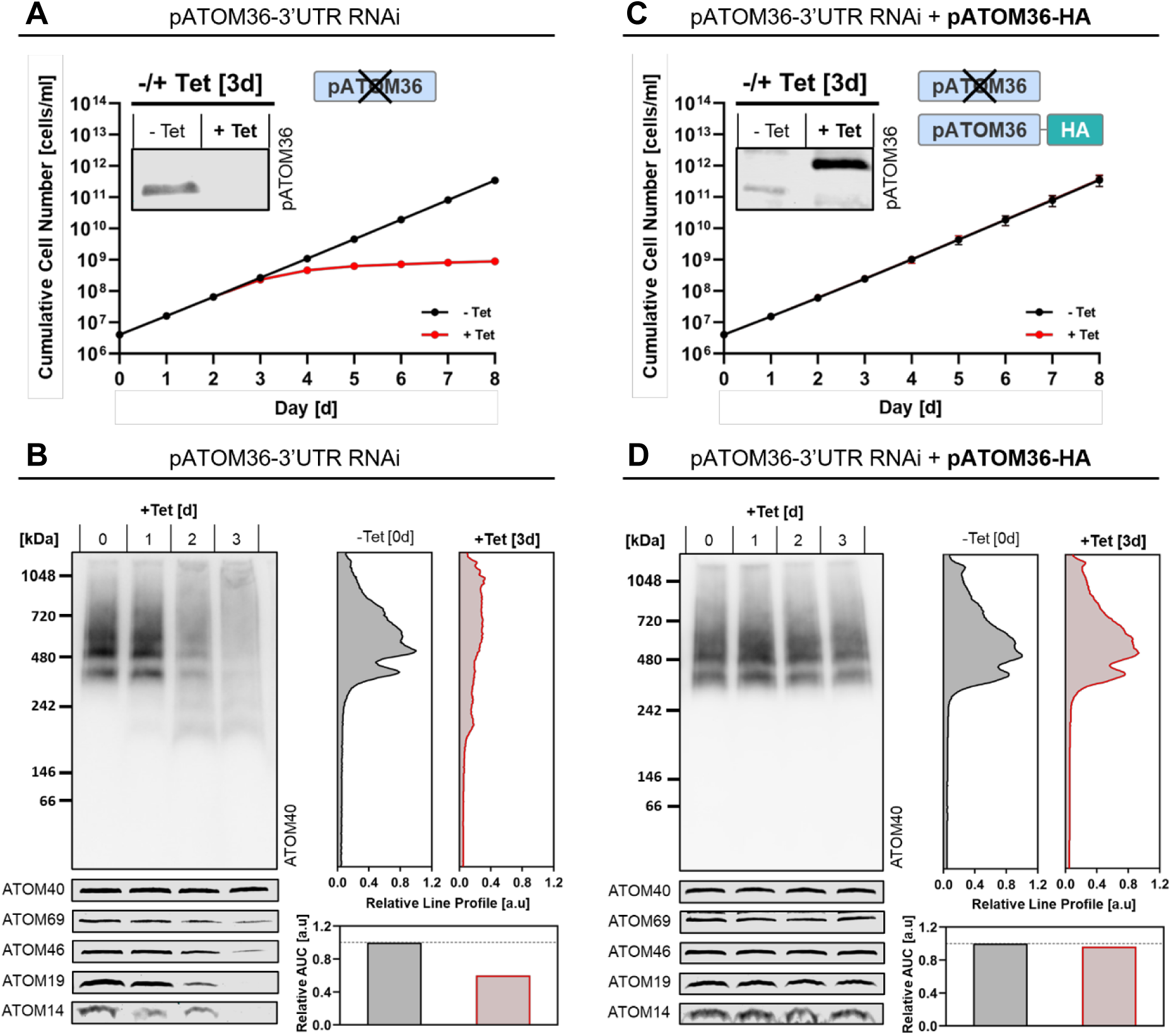
*In vivo* pATOM36-3ʹUTR RNAi system in procyclic *T. brucei* and analysis of ATOM complex assembly and subunit abundance. Growth analysis of procyclic *T. brucei* cell lines under tetracycline-induced (+Tet) and non-induced (−Tet) conditions. pATOM36 knockdown efficiency and expression of C-terminally 3×HA-tagged wild-type pATOM36 (pATOM36-HA) were confirmed by immunoblotting using a pATOM36-specific antibody. Mitochondrial membrane extracts from the indicated cell lines were analyzed for ATOM complex assembly over three days of +Tet induction by Blue Native PAGE, followed by immunoblotting with an ATOM40-specific antibody. Relative intensity profiles of ATOM40-containing complexes from -Tet and day three +Tet samples were quantified in ImageJ, and the relative area under the curve (AUC) was calculated. The same extracts were immunoblotted for ATOM40 and the putative pATOM36 substrates ATOM69, ATOM46, ATOM19, and ATOM14. **A)** Growth and RNAi evaluation of the parental pATOM36-3ʹUTR RNAi cell line. **B)** Analysis of ATOM complex assembly and abundance of putative pATOM36 substrates in the parental pATOM36-3ʹUTR RNAi cell line. **C)** Growth analysis and validation of RNAi knockdown and protein expression in the pATOM36-3ʹUTR RNAi + pATOM36-HA cell line. **D)** Analysis of ATOM complex assembly and abundance of putative pATOM36 substrates in the pATOM36-3ʹUTR RNAi + pATOM36-HA cell line.

Expression of the C-terminally 3×HA-tagged wild-type pATOM36 (pATOM36-HA) in the pATOM36-3ʹUTR RNAi cell line restores growth to wild-type levels, despite almost complete depletion of endogenous pATOM36 (Fig. 2C). Moreover, not only growth but also the ATOM complex assembly defect and the steady-state levels of all tested subunits were reconstituted to wild-type levels (Fig. 2D). These experiments showed that C-terminally 3xHA-tagged pATOM36 is fully functional.

### Untagged MTCH1 replaces pATOM36 function in OMM protein biogenesis

Next, we used the *in vivo* system described above to assess whether human MTCH1 or MTCH2 can replace the OMM protein biogenesis function of pATOM36. To detect MTCH1 and MTCH2 in transgenic trypanosomes, we added a 3×HA tag to their C-termini, and the two tagged proteins were individually expressed in the pATOM36-3’UTR RNAi cell line. The results showed that neither MTCH1-HA (Fig. 3A, left) nor MTCH2-HA (Fig. 3B, left) could rescue the growth arrest caused by depletion of endogenous pATOM36. Moreover, digitonin-based cell fractionation revealed that only small quantities of MTCH1-HA (Fig. 3A, right) and even lower amounts of MTCH2-HA (Fig. 3B, right) co-fractionated with the crude mitochondrial fraction. Finally, BN-PAGE analyses showed that, as expected, ATOM complex assembly was still disrupted in both the MTCH1-HA (Fig. 3C) and MTCH2-HA (Fig. 3D) cell lines.

**Figure 3:**
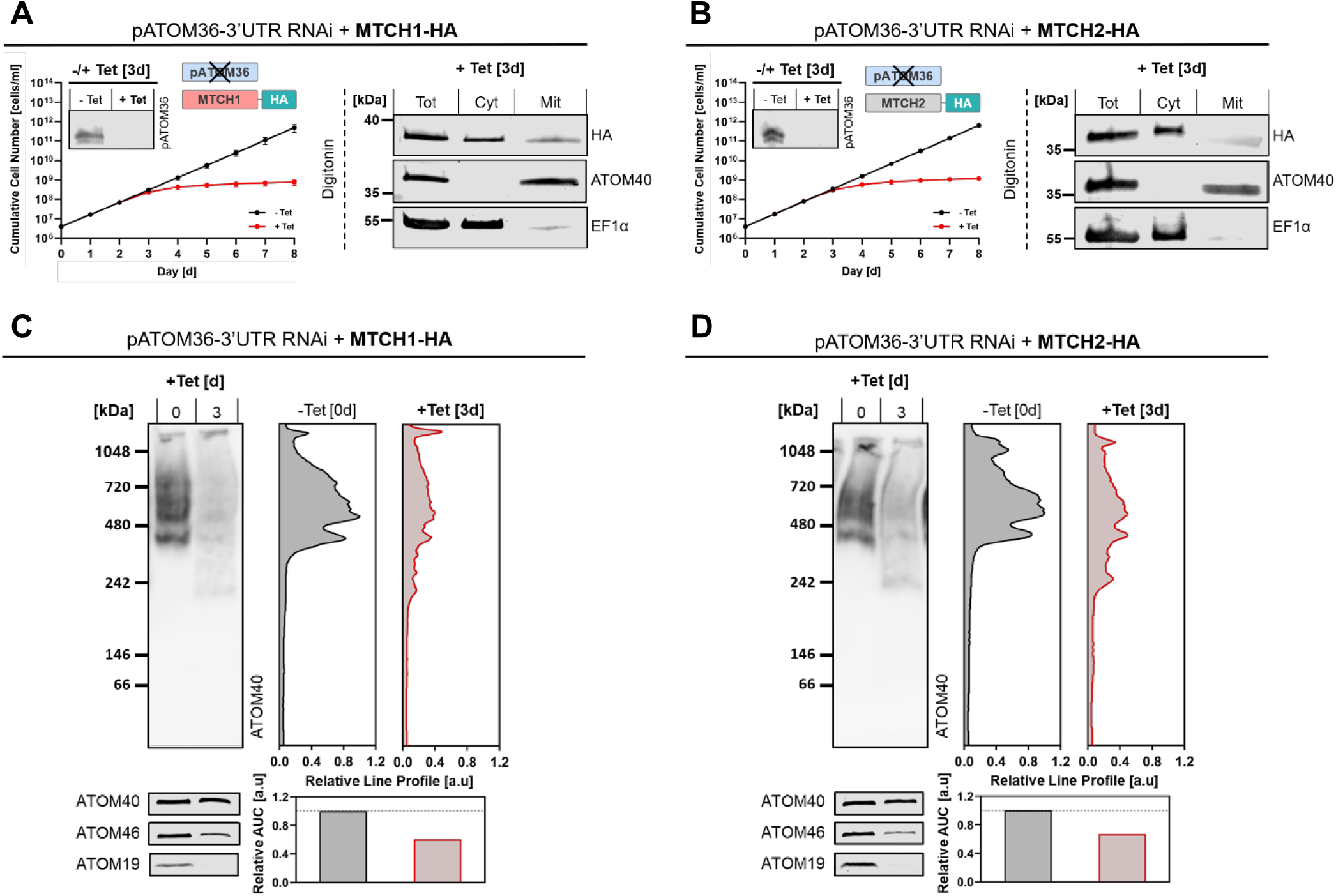
Heterologous expression of HA-tagged MTCH1/2 in pATOM36-3ʹUTR RNAi procyclic *T. brucei* and analysis of mitochondrial localization and ATOM complex integrity. Growth of individual cell lines was evaluated under +Tet and -Tet conditions. pATOM36 knockdown efficiency was verified by immunoblotting using a pATOM36-specific antibody. Expression of MTCH1 and MTCH2 constructs and their co-fractionation with mitochondria were assessed at day three +Tet by immunoblot analysis of total (Tot), digitonin-extracted cytosolic (Cyt), and mitochondrial (Mit) fractions from cell lines expressing C-terminally 3×HA-tagged MTCH1 (MTCH1-HA) or MTCH2 (MTCH2-HA). ATOM40 F1α served as mitochondrial and cytosolic markers, respectively. Mitochondrial membrane extracts were analyzed for ATOM complex assembly after three days of +Tet induction by BN-PAGE followed by ATOM40 immunoblotting. Relative intensity profiles of ATOM40-ining complexes from −Tet and day three +Tet samples were quantified in ImageJ, and the relative AUC was calculated subsequently. same extracts were immunoblotted for ATOM40 and the putative substrates ATOM46, and ATOM19. Growth and RNAi evaluation, ed by digitonin fractionation, in **A)** the pATOM36-3ʹUTR RNAi + MTCH1-HA cell line and **B)** the pATOM36-3ʹUTR RNAi + MTCH2-HA cell analysis of ATOM complex assembly and abundance of putative pATOM36 substrates in **C)** the pATOM36-3ʹUTR RNAi + MTCH1-HA cell nd **D)** the pATOM36-3ʹUTR RNAi + MTCH2-HA cell line.

To determine whether the C-terminal 3×HA tag interfered with mitochondrial localization of MTCH1 or MTCH2, the experiment described above was repeated using wild-type versions of the two proteins. The result for untagged MTCH1 showed that its expression could partially complement the growth defect (Fig. 4A, left). Moreover, using an antibody recognizing untagged MTCH1, immunoblots of a digitonin-based cell fractionation demonstrated that MTCH1 was now exclusively detected in the crude mitochondrial fraction (Fig. 4A, right).

**Figure 4:**
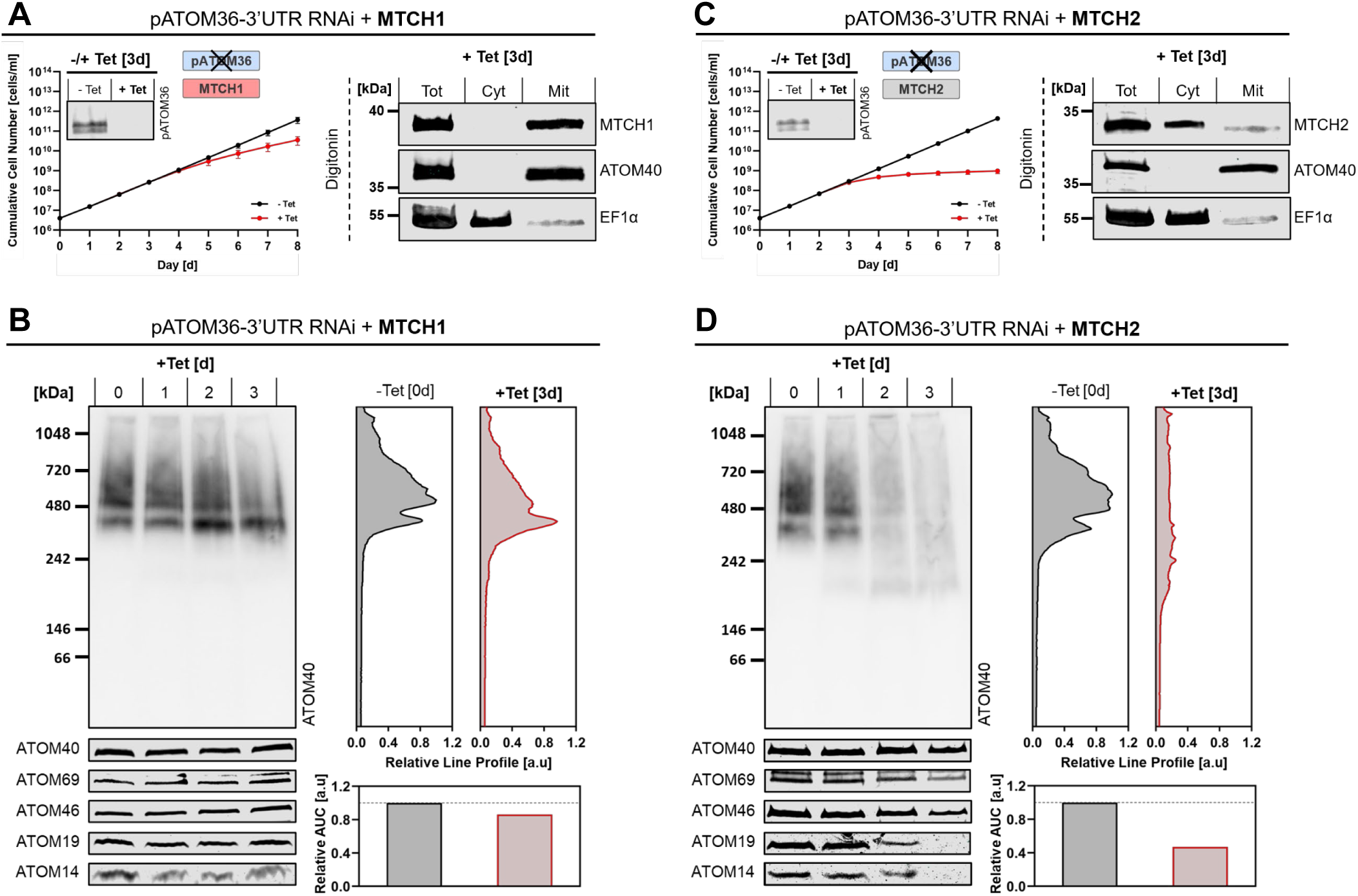
Heterologous expression of native MTCH1 and MTCH2 in pATOM36-3ʹUTR RNAi procyclic *T. brucei* and analysis of mitochondrial localization and ATOM complex integrity. Growth of cell lines was assessed under +Tet and −Tet conditions, and pATOM36 knockdown was med by immunoblotting with a pATOM36-specific antibody. Expression and mitochondrial co-fractionation of untagged MTCH1 and 2 were analyzed at day three +Tet by immunoblotting total (Tot), digitonin-extracted cytosolic (Cyt), and crude mitochondrial (Mit) fractions. ATOM40 and EF1α served as mitochondrial and cytosolic markers, respectively. Mitochondrial membrane extracts were analyzed OM complex assembly over the three-day + Tet induction by BN-PAGE and ATOM40 immunoblotting. Relative intensity profiles of 40-containing complexes from −Tet and day three +Tet samples were quantified in ImageJ, and the relative AUC was calculated. The extracts were immunoblotted for ATOM40 and the putative substrates ATOM69, ATOM46, ATOM19, and ATOM14. Characterization of M36-3ʹUTR RNAi + MTCH1 cells in **A)** growth, RNAi validation, and crude mitochondrial co-fractionation and **B)** ATOM complex assembly abundance of putative substrates. Evaluation of the pATOM36-3ʹUTR RNAi + MTCH2 cell line in **C)** growth, RNAi validation, and crude mitochondrial co-fractionation and **D)** ATOM complex assembly and abundance of putative substrates.

Thus, in *T. brucei*, unlike in *S. cerevisiae*^16^, the C-terminal 3xHA tag interfered with mitochondrial localization of MTCH1. Most importantly, BN-PAGE analysis revealed that untagged MTCH1 could restore ATOM complex assembly (Fig. 4B). The only qualitative difference detected in the ATOM complex pattern after complementation was a small shift towards the lower of the two main ATOM40-containing complexes. Moreover, the steady-state levels of all tested ATOM subunits were also restored to wild-type levels. These results demonstrated that mitochondria localized MTCH1 could replace the OMM biogenesis function of pATOM36. Note that the inability of MTCH1 to completely restore the growth phenotype is expected, as it cannot replace the TAC function of pATOM36.

Untagged MTCH2, in contrast, did not complement the growth defect (Fig. 4C, left), and immunoblot analysis of a digitonin-based cell fractionation using an antibody recognizing untagged MTCH2 showed only minimal quantities of the protein in the crude mitochondrial fraction (Fig. 4C, right). Consequently, neither restoration of ATOM complex assembly on BN-PAGE nor restoration of the steady-state levels of the tested ATOM subunits were observed. Thus, the results for the untagged MTCH2 closely mirrored those of the C-terminally 3×HA-tagged version of the protein (Fig. 4D).

### A MTCH1-MTCH2 chimera compensates for the lack of pATOM36

Even though human MTCH1 and MTCH2 are both OMM proteins^17,31^, only MTCH1 was efficiently targeted to the mitochondrion when expressed in *T. brucei* (Fig. 4A). An alignment of the two proteins shows a sequence identity of 48%, including the typical repeat structure of the SLC25 family of mitochondrial carrier proteins (Fig. 5A). The main difference between the two is that MTCH1 has an N-terminal extension of 78 aa that is absent from MTCH2 and any other mitochondrial protein (Fig. 5A). The extension is particularly rich in alanine, glycine, arginine, and proline, has a net positive charge, and according to the InterScanPro algorithm, likely represents an intrinsically disordered region^32^.

**Figure 5:**
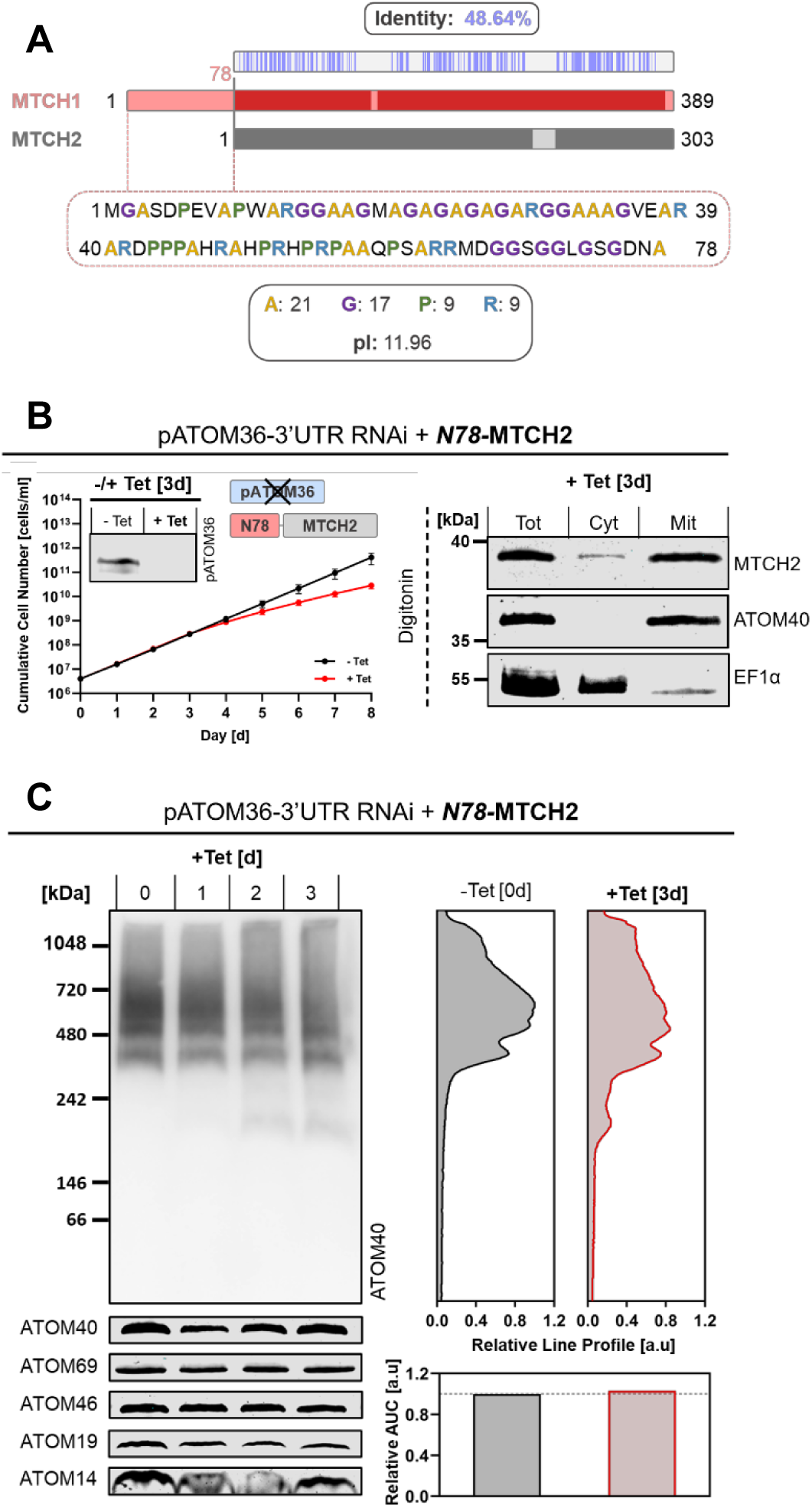
Sequence comparison of MTCH1 and MTCH2 reveals a unique N-terminal extension in MTCH1. **A)** Sequence alignment of MTCH1 and MTCH2 using UniProt Align shows strong conservation in the core regions (∼49% identity), with MTCH1 containing a unique N-terminal extension (residues 1–78) absent in MTCH2. The amino acid composition of this extension is shown graphically, highlighting alanine (orange), glycine (violet), proline (green), and arginine (blue). This N-terminal region is enriched in basic residues and predicted to carry a net positive charge at neutral pH. **B)** Growth and RNAi validation of pATOM36-3ʹUTR RNAi procyclic *T. brucei* cells expressing the N78-MTCH1–MTCH2 fusion protein (N78-MTCH2). Subcellular localization was assessed by immunoblotting of total (Tot), digitonin-extracted cytosolic (Cyt), and crude mitochondrial (Mit) fractions, with ATOM40 and EF1α as mitochondrial and cytosolic markers, respectively. **C)** ATOM complex assembly in mitochondrial extracts from the N78-MTCH2 cell line was monitored over three days of +Tet by BN-PAGE and ATOM40 immunoblotting. ATOM40-containing complexes were quantified in ImageJ to compare relative AUC values, and the same extracts were probed for ATOM40, ATOM69, ATOM46, ATOM19, and ATOM14.

Thus, we tested whether expression of a chimeric protein (N78-MTCH2), consisting of the N-terminal 78 aa extension of MTCH1 fused to full-length MTCH2, could complement for the depleted endogenous pATOM36 in the pATOM36-3’UTR RNAi cell line. This was indeed the case, N78-MTCH2 was efficiently targeted to the mitochondrion and could partially restore growth retardation of the RNAi cell line (Fig. 5B) to the same extent as MTCH1 (Fig. 4A). Most importantly, ATOM complex formation and steady-state levels of the tested ATOM subunits were restored to wild-type levels (Fig. 5C). The only difference between N78-MTCH2 and MTCH1 evident in the BN-PAGE analysis was the appearance of a small amount of an approximately 200 kDa ATOM40-containing subcomplex that was absent in the MTCH1 complementation (Fig. 4B).

In summary, we conclude: i) that the N-terminal 78 aa extension of MTCH1 allowed efficient mitochondrial targeting of MTCH2, and ii) that either MTCH1 or N78-MTCH2 can replace the OMM protein biogenesis function of pATOM36.

### MTCH1 and N78-MTCH2 cannot restore kDNA segregation defects

pATOM36 functions in the biogenesis of α-helical OMM proteins, including many ATOM subunits. It has been described in a variety of organisms that perturbation of mitochondrial protein import can change mitochondrial morphology^6,33^. This was also observed after depletion of pATOM36, which caused the normally extended network-like mitochondrion of procyclic *T. brucei* to adopt a more condensed structure (Fig. 6A, lower left panel)^26,27^. Expression of full-length pATOM36-HA, MTCH1 or N78-MTCH2 in the pATOM36-3’UTR cell line restored normal mitochondrial morphology (Fig. 6A) in line with the observed ATOM complex assembly. The same had previously been observed after co-expression of the yeast Mim1 and Mim2^30^.

**Figure 6:**
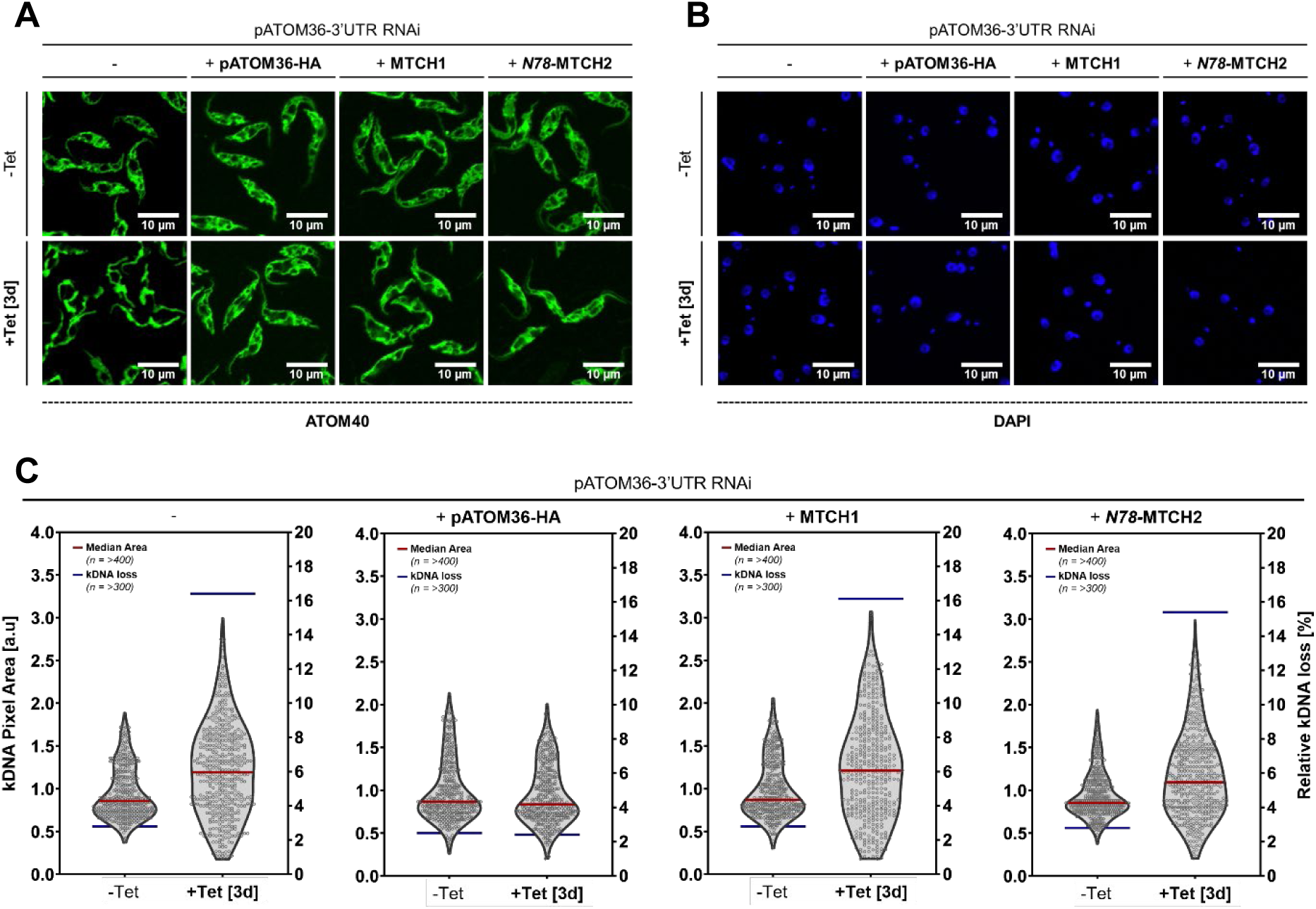
Analysis of mitochondrial morphology and kDNA segregation in procyclic *T. brucei* cells expressing MTCH1 and N78-MTCH2. Immunofluorescence microscopy of the different -Tet and +Tet procyclic *T. brucei* cell lines. **A)** Mitochondrial marker ATOM40 (green). **B)** Nuclear and kinetoplast DNA stained with DAPI (blue). **C)** kDNA segregation was evaluated by measuring kDNA area in −Tet and three-day +Tet cells. DAPI-stained fluorescence images were analyzed using a semi-automated ImageJ macro to quantify the pixel area of individual kDNAs, with the red line indicating the median kDNA area (n = >400). The kDNA loss phenotype, associated with segregation defects, was assessed manually by overlaying DAPI fluorescence images onto corresponding brightfield images, with the blue line representing the relative extent of kDNA loss (n = >300).

In addition to its role in the biogenesis of OMM proteins, pATOM36 is also an essential subunit of the TAC, where it ensures proper segregation of the replicated kDNA. Depletion of TAC subunits, including pATOM36^26^ cause kDNA segregation defects but does not interfere with kDNA replication^34^. This results in over-replicated kDNAs in some cells and complete loss of kDNAs in other cells. Moreover, due to asymmetric distribution of the replicated kDNAs, cells having very small kDNAs are also observed. All these outcomes will eventually lead to cell death.

The amount of kDNA in a given cell can be determined by DAPI-staining (Fig. 6B) and subsequent quantification of the fluorescent signal (Fig. 6C). Analysis of the kDNA contents of the pATOM36-3’UTR RNA cell line showed that depletion of the protein caused kDNA loss in many cells (Figure 6C, blue bar). Moreover, in the cells that still contained kDNA, it was either largely over-replicated or much smaller than in uninduced cells (Fig. 6C, left panel).

Whereas expression of full-length pATOM36-HA restored kDNA segregation, this defect could not be rescued by expression of MTCH1 or N78-MTCH2. This is expected because no structure comparable to the TAC is found in human cells, MTCH1 or N78-MTCH2 can therefore not replace the role that pATOM36 plays as a TAC subunit.

### pATOM36 facilitates bidirectional lipid transport in liposomes

It has recently been shown that membrane protein insertases such as YidC, Oxa1, and MTCH2 exhibit lipid-scrambling activity, i.e. non-directed enzyme-catalyzed lipid exchange between the two bilayer leaflets^35^. To test whether pATOM36 also exhibits similar activity, we expressed and purified the protein in *S. cerevisiae*. For this purpose, we used a fusion construct in which pATOM36 was linked to yeast-enhanced green fluorescent protein via a linker containing a Tobacco Etch Virus (TEV) protease cleavage site. The fusion protein (pATOM36–GFP) carried a C-terminal 8×His tag, allowing efficient purification (Fig. 7A).

**Figure 7:**
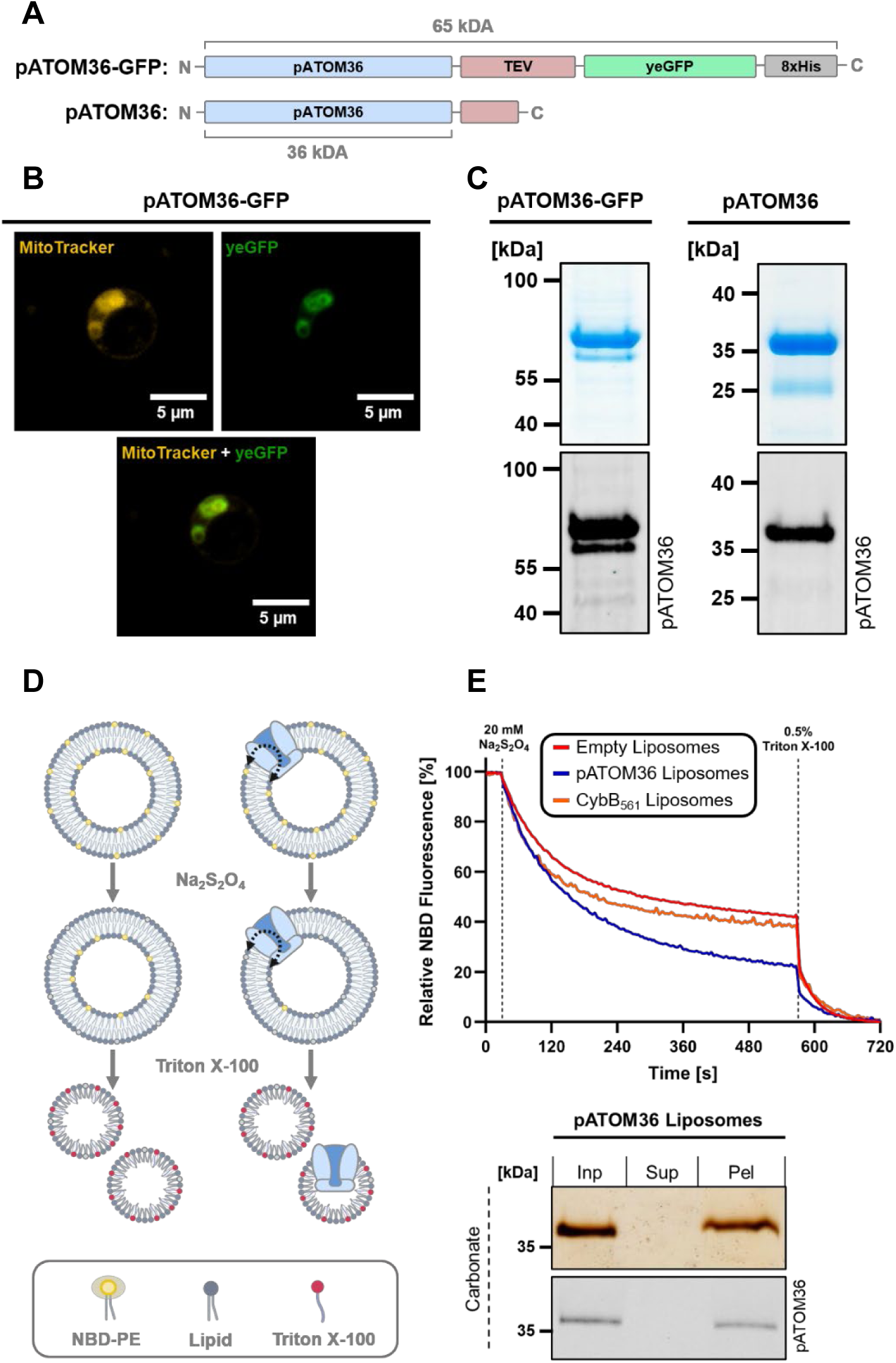
pATOM36 expressed in *S. cerevisiae*, purified in detergent, and reconstituted into liposomes exhibits lipid scrambling activity *in vitro*. **A)** Schematic representation of the pATOM36-GFP construct used for overexpression and the TEV-digested pATOM36 obtained after the purification procedure. **B)** Immunofluorescence microscopy of pATOM36-GFP (green) in yeast, co-stained with MitoTracker (yellow) to visualize mitochondria. **C)** SDS-PAGE and immunoblot of non-digested pATOM36-GFP and TEV-digested pATOM36 following detergent solubilization and Ni-NTA purification. **D)** Schematic of the *in vitro* lipid-scrambling assay. pATOM36 is light blue, lipids gray, fluorescent NBD-PE yellow, and Triton X-100 red. **E)** Time-course fluorescence measurements of lipid scrambling in pATOM36 proteoliposomes (blue) versus empty liposomes (red) and CybB561 proteoliposomes (orange). Liposomal membrane integration of pATOM36 was further assessed by alkaline carbonate extraction, where reconstituted proteins sediment in the pellet (Pel) and non-reconstituted proteins remain in the supernatant (Sup), compared to total input (Inp) by SDS-PAGE and immunoblotting.

Fluorescence microscopy combined with MitoTracker staining showed that pATOM36-GFP was efficiently localized to yeast mitochondria (Fig. 7B). Detergent solubilization followed by Ni–NTA affinity purification and TEV protease cleavage allowed the isolation of both the pATOM36–GFP fusion protein and the cleaved pATOM36. (Fig. 7C). Subsequently, scramblase activity was measured using an established in vitro assay as described (Fig. 7D)^36^. Cleaved pATOM36 was reconstituted into preformed liposomes containing 0.5 % fluorescent lipids (nitrobenzoxadiazole phosphoethanolamine, NBD-PE), while liposomes containing no protein served as control. Addition of the non-membrane permeable reducing agent sodium dithionite leads to irreversible fluorescence quenching by chemical reduction of the NBD fluorophores in the outer, but not the inner bilayer leaflet. In the absence of scramblase activity, this reduction in fluorescence is expected to be slightly above 50% and thus a residual fluorescence of <50%, as the outer leaflet contains more lipids due to the increased radius. In the presence of scramblase activity, however, a higher reduction of fluorescence is expected at later time points as lipids from the inner leaflet flip to the outer leaflet, where they react with sodium dithionite and their fluorescence is quenched. Subsequent addition of detergent disrupts the liposomes, allowing sodium dithionite to also react with the NBD fluorophores of the inner leaflet, yielding 100% fluorescence quenching ^36^. The results of our experiments are depicted in Fig. 7E. In the control experiment with empty liposomes, where no lipid exchange between leaflets can take place, addition of sodium dithionite and detergent lead to a remaining fluorescence value of ∼45%, in good agreement with the expectation that the outer leaflet is slightly larger. Importantly, in the liposomes containing TEV-processed pATOM36, residual fluorescence was only ∼20% upon sodium dithionite addition, indicating enzyme-mediated lipid exchange between the two leaflets. To confirm that the observed activity is a specific feature of pATOM36 and not of any integral membrane protein, the same experiment was performed with liposomes containing *E. coli* cytochrome *b561* (CybB_561_), which has four transmembrane helices and has not been associated with lipid scrambling activity^37^. CybB_561_-containing liposomes had almost identical NBD quenching activity to empty liposomes (Fig. 7E). Thus, we conclude that pATOM36, as many protein insertases, has scramblase activity, and thus facilitates bidirectional lipid transport.

### pATOM36 and MTCH1/MTCH2 form intra-membranous cavities

Given the analogous function of pATOM36 and MTCH1/MTCH2, we compared their predicted structures using AlphaFold3^38^ and simulated how they are inserted into the OMM using the PPM 3.0 algorithm^39^. Models of both proteins were of high-quality, with very good pTM confidence scores in the respective TMDs (Sup. Fig. 1)

The pATOM36 structure comprised five TMDs with its C- and N-terminus exposed to the cytosol and IMS, respectively, in accordance with previous experiments (Fig. 8A, top)^25^. The hydrophobic core thickness of a typical lipid bilayer is ∼28–35 Å, with the OMM at the lower end of this range, reflecting its low content of membrane-thickening cholesterol and sphingolipids^40^. Our model indicates that pATOM36 further reduces the hydrophobic core thickness of the OMM to ∼23.7 Å. In contrast, analysis of the ATOM subunit ATOM19, predicted to have two α-helical TMDs yielded a value of 27.7 Å (Sup. Fig. 2). In the model, TMD1 of pATOM36 adopts a discontinuous α-helix, whereas TMD2–TMD5 form continuous membrane-spanning α-helices (Fig. 8A, bottom). TMD4 and TMD5 contain GxxxG motifs, which might facilitate close helix-helix interactions, reflected by their spatial proximity within the model^41^. The five TMDs in the pATOM36 model form a polar cavity containing charged residues. The cavity is ∼24 Å wide, extends ∼22 Å into the lipid-bilayer, and is open towards the cytosol (Fig. 8B). Towards the IMS side, it is sealed at the level of the membrane border by close interactions of TMDs 3 to 5 and additional protein elements. TMDs 2 to4 are strongly amphipathic. While the surface facing the lipid bilayer is strongly hydrophobic, the helices contain many positively and few negatively charged amino acids towards the cavity, strongly suggesting that it is in a direct aqueous continuum with the cytoplasm. This strongly hydrophilic surface forms the back wall of a U-shaped cavity that becomes progressively less polar towards its lateral opening to the hydrophobic membrane core (Fig. 8C). The sides towards the lateral opening are defined by TMD1 and TMD5. The cavity is completed by a hydrophobic rim at the lateral opening, allowing contact to the hydrophobic fatty acids tails of the lipid bilayer (Fig. 8C).

**Figure 8:**
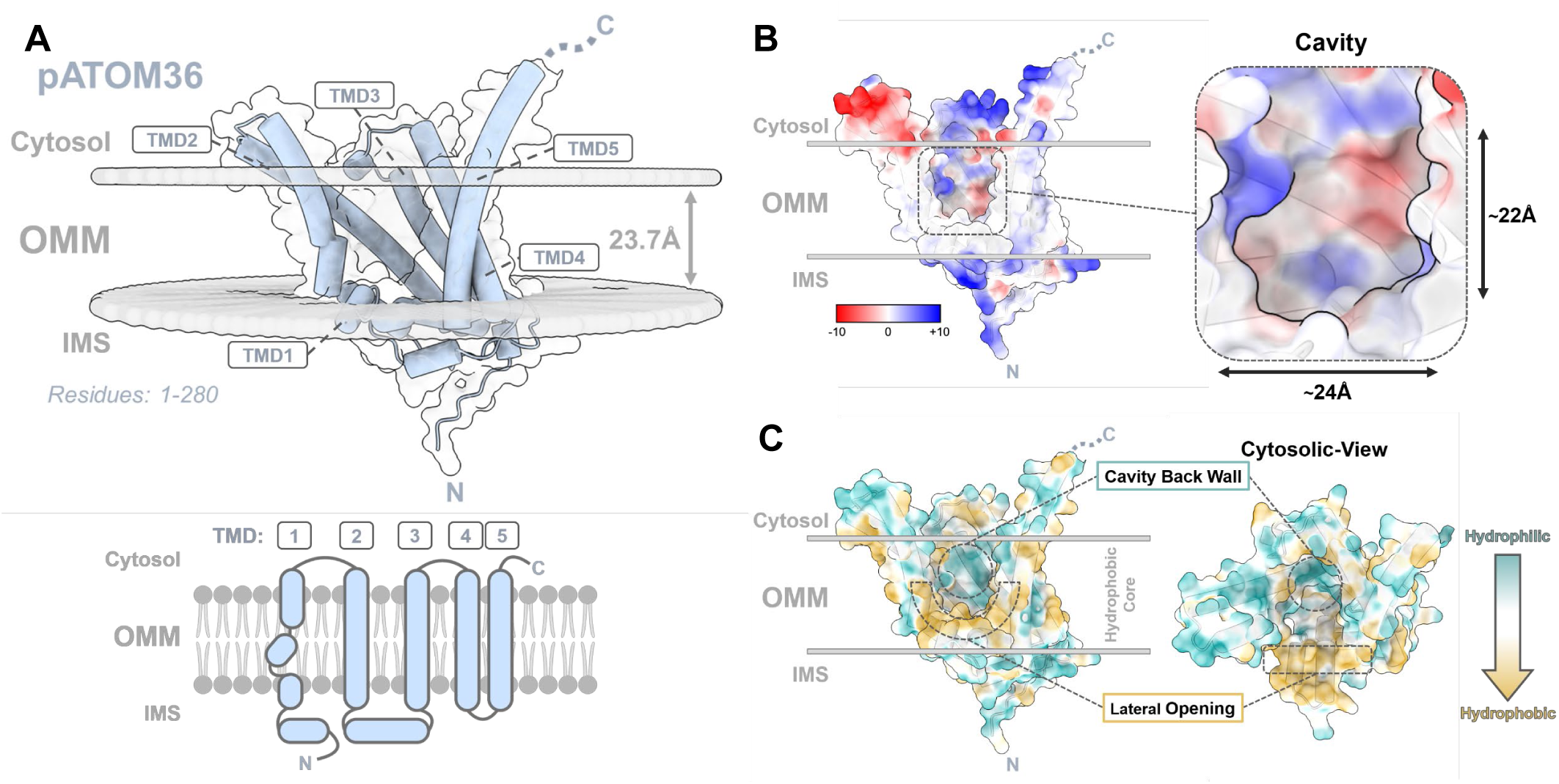
Structural insights into *T. brucei* pATOM36. AlphaFold3 was employed to generate a structural model of pATOM36 (light blue), was computationally inserted into the OMM using the PPM 3.0 server. **A)** *Top:* **T**he pATOM36 model (residues 1–280) showing five predicted transmembrane domains (TMDs), embedded within the OMM (grey). Cartoon representation with the hydrophobic membrane reduced to 23.7 Å (grey arrow), C-terminus facing the cytosol, and N-terminus toward the IMS. *Bottom:* Schematic sketch of the 5 predicted TMDs, comprising four continuous and one discontinuous helix. **B)** Electrostatic surface representation highlighting positively (blue) negatively (red) charged residues within a cavity that extends ∼22 Å into the hydrophobic membrane core and is ∼24 Å wide. **C)** Hydrophobicity surface representation showing hydrophobic (brown) and hydrophilic (cyan) regions. Dashed circles mark the back wall of the cytosol-accessible cavity and the U-shaped rim of the lateral gate facing lipid alkyl chains, both located within the hydrophobic membrane Cytosolic view reveals an amphipathic cavity architecture, characterized by hydrophilic back walls transitioning into hydrophobic lateral gates.

Genetically, MTCH1 belongs to the SLC25 family of carrier proteins, typically located exclusively in the inner mitochondrial membrane, where they are crucial for transport of mostly negatively charged compounds like ADP, ATP, phosphate, pyruvate and others between cytoplasm and mitochondrial matrix^42^. They form bundles consisting of six TMDs organized into three homologous repeats (∼100 amino acids each) where both termini face the IMS. Each of the repeats consists of a discontinuous α-helix with the characteristic Px[DE]xx[KR] motif, where a kink is introduced by a conserved proline, followed by a short IMS-facing loop and a continuous α-helix^43,44^.

Next to SLC25A46, a protein involved in mitochondrial lipid homeostasis^45^, MTCH1 and MTCH2 are to date the only reported members of the SLC25 family to be localized in the OMM^15^. Some differences are found if compared to a native family member such as the ADP/ATP translocase^42,44^. In MTCH, the Px[DE]xx[KR] motif is only partially conserved in the odd-numbered TMDs 1, 3, and 5, in some cases retaining only the proline residue^15^. Importantly, the structural prediction resolves only five TMDs (Fig. 9A, top), which superimpose well with TMD1 to 5 of the ADP/ATP (SLC25A4) translocase (Sup. Fig. 3). The absence of a canonical TMD6 in MTCH1 is supported by InterProScan motif annotations^32^ and consistent with trypsin digestion experiments with Presenilin-associated protein (earlier name of MTCH1) that showed that its C-terminus reaches into the IMS (Fig. 9A, bottom)^46^.

**Figure 9:**
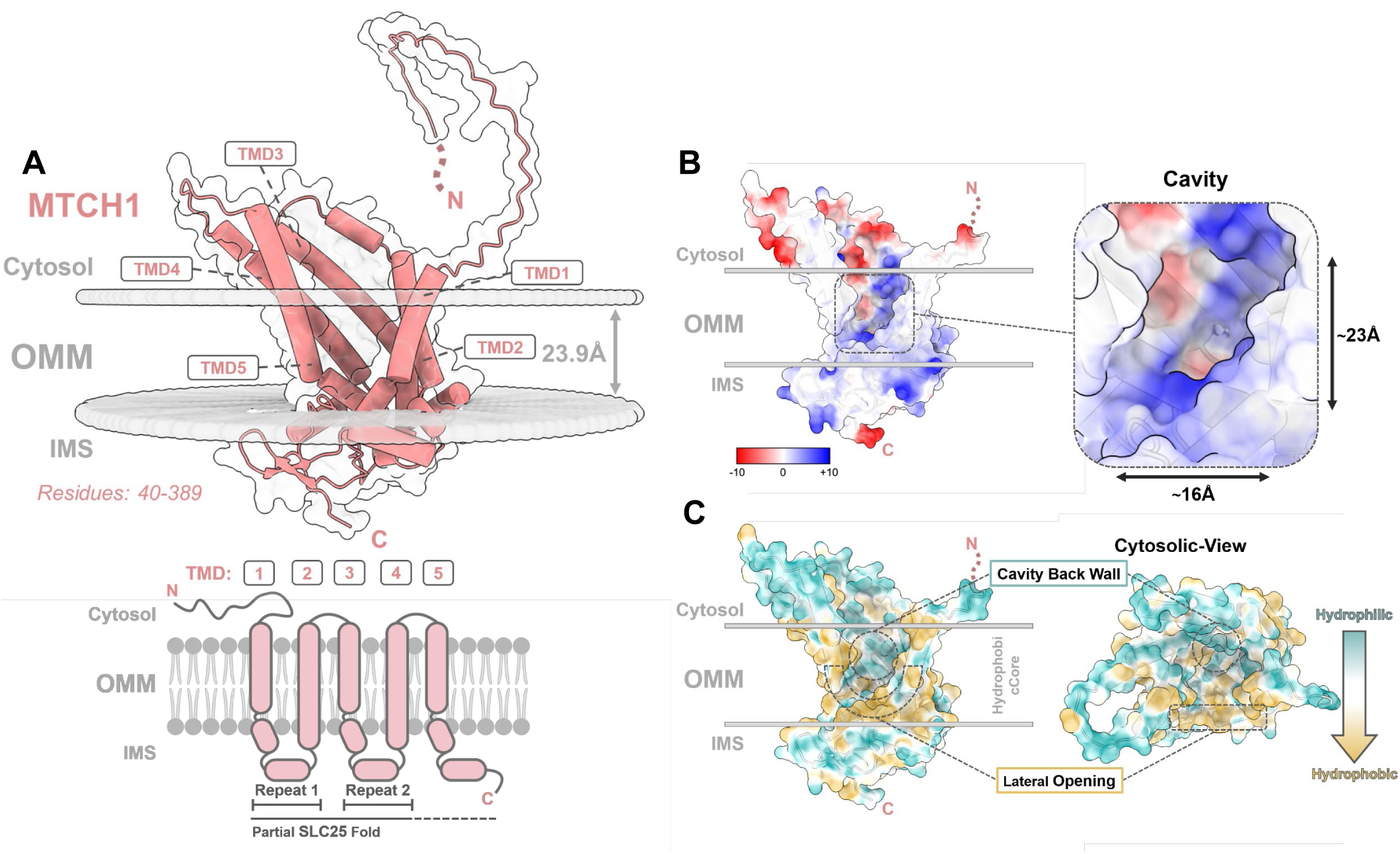
Predicted structural features of human MTCH1. The AlphaFold3 model of MTCH1 (coral) was generated and computationally added in the OMM using the PPM 3.0 server. **A)** *Top:* Cartoon representation of the MTCH1 model (residues 40-389) within the OMM, revealing five predicted TMDs and an extension at the N-terminus. The hydrophobic membrane core is reduced to 23.9 Å (grey arrow), he C-terminus facing the IMS and the N-terminus oriented towards the cytosol. *Bottom:* Schematic sketch of the five predicted TMDs, partially following the SLC25 fold. **B)** Electrostatic surface mapping highlights positively (blue) and negatively (red) charged residues lining a that extends ∼23 Å into the hydrophobic membrane core and spans ∼16 Å in width. **C)** Hydrophobicity surface representation shows phobic (brown) and hydrophilic (cyan) regions. Dashed circles mark the position of the back wall of the cavity and the U-shaped rim of lateral gate. Similar to pATOM36, the cytosolic view shows an amphipathic cavity architecture with a hydrophilic back wall and phobic lateral gate.

The lack of TMD6 leads to a physical gap within the helical bundle and provides a structural solution for the necessity of a lateral opening towards the membrane. In addition, the charges found within canonical members of the SLC25 family to bind charged metabolites, are an ideal template to build a hydrophilic cavity. Like pATOM36, calculations predict that MTCH1 also reduces the hydrophobic thickness of the OMM (to ∼23.9 Å). The structural model of MTCH1 predicts that the TMDs 1-5 form a hydrophilic cavity (∼23 Å deep and ∼16 Å wide) that is accessible from the cytosol and laterally open toward the hydrophobic membrane core (Fig. 9B). Similar to pATOM36, the back wall of the MTCH1 cavity is formed by amphipathic α-helices TMDs2-TMD4 carrying their polar and charged residues towards the inside of the cavity. The IMS-exposed loops of the three repeats seal the cavity on the IMS side (Fig. 9B).

The hydrophobicity maps of MTCH1 show a similar distribution of hydrophilic and hydrophobic regions within the cavity with a pronounced hydrophobic U-shaped rim defining the contact site with the hydrophobic core of the membrane (Fig. 9C)

Overall, the structure of both MTCH1 is very similar to the one of pATOM36, both in overall architecture as well as its local surface properties.

The predicted model for MTCH2 is essentially identical to MTCH1, but the orientation of MTCH2 in the OMM has not yet been experimentally determined (Sup. Fig. 4).

### Conserved cavity structure despite different topologies

As described in the paragraph above, MTCH1 and pATOM36 display a striking overall resemblance, i.e. hydrophilic cavity with access towards the cytosol, despite showing a very low sequence identity of 22% (MTCH2, 19%), suggesting extensive evolutionary divergence. While both proteins form the hydrophilic cavity using five transmembrane domains, their membrane orientation (topology) is inverted. MTCH1 has an N_Cyt_ and C_IMS_ orientation, whereas the topology of pATOM36 is N_IMS_ and C_Cyt_, likely contributing to the low sequence identity.

To compare similarity at the structural level, we used TopMatch, which permits permutation (or shuffling) of individual secondary-structure elements in three-dimensional space while preserving their internal N-to-C directionality^47^. The best such alignment of pATOM36 and MTCH1 preserves the core topology of both proteins very well (Fig. 10A). The structural similarity is confined to TMDs1 to TMD4 of the two proteins, whereas TMD5 did not align well. Consistent with the inverted topology, odd TMDs now align with even numbered TMDs, i.e. TMD1_pATOM36_ aligns with TMD4_MTCH1_, TMD2_pATOM36_ with TMD3_MTCH1_, TMD3_pATOM36_ with TMD2_MTCH1_, and TMD4_pATOM36_ with TMD1_MTCH1_. In this alignment, the N-to-C direction of corresponding transmembrane helices is preserved in both proteins; however, the shuffling of the helices results in different loop connectivities (Fig. 10B). For the alignment with pATOM36, TopMatch predicted three permutations in MTCH1 (resulting in 4 segments), leading to the identification of 122 structurally equivalent amino acid pairs (Fig. 10C, Sup. Fig. 5).

**Figure 10:**
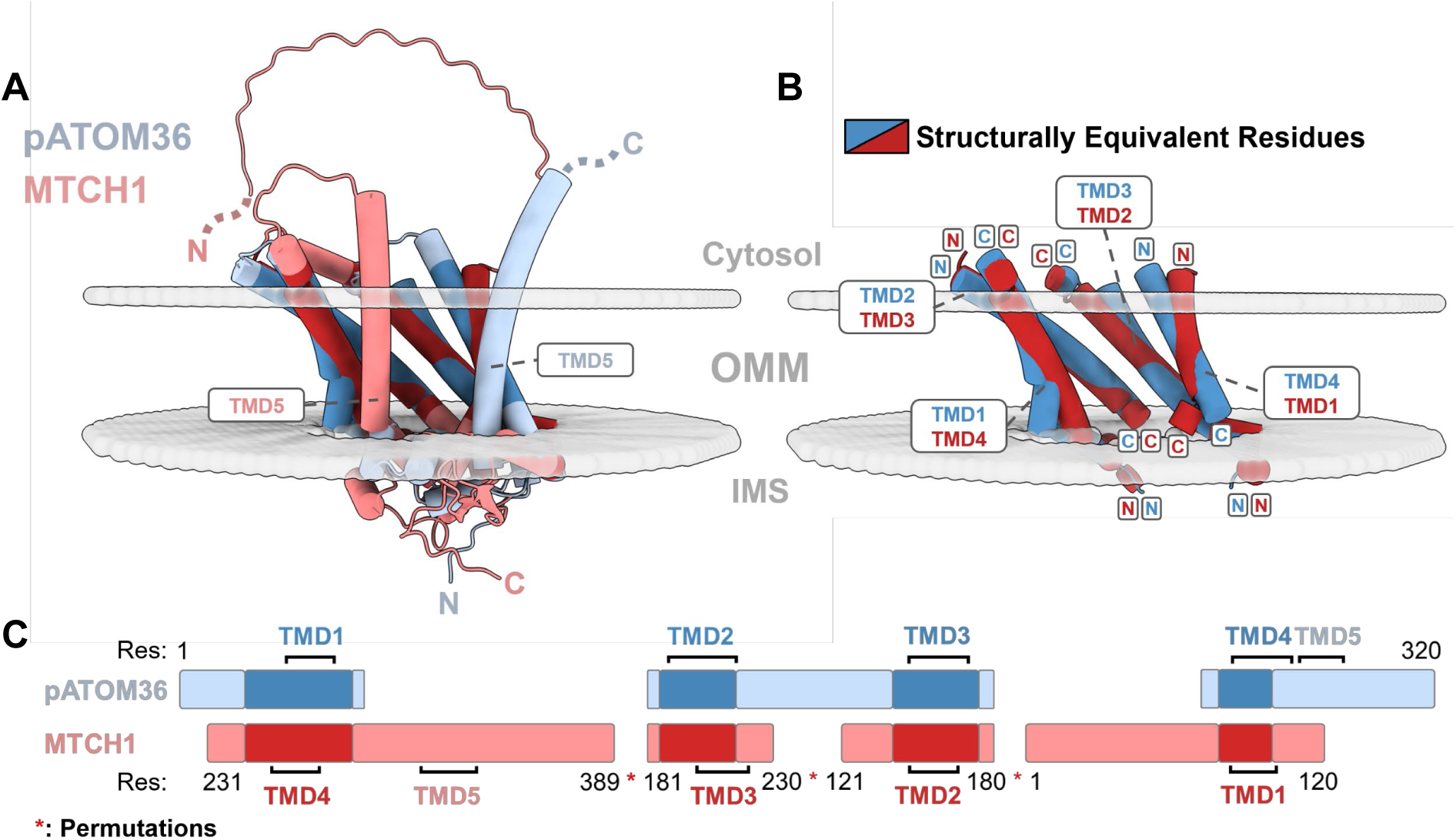
Pairwise structural alignment reveals a conserved cavity back wall architecture in pATOM36 and MTCH1. The Pairwise structural alignment of pATOM36 (light blue) with MTCH1 (coral) was performed using the TopMatch algorithm. **A)** Global structural alignment shows that pATOM36 and MTCH1 adopt inverted topologies that nevertheless enable cytosol-accessible cavities in both insertases. **B)** Structurally equivalent residues between pATOM36 (dark blue) and MTCH1 (dark red) are mainly confined to the TMDs, with each segment maintaining identical local topology. **C)** Schematic representation of the structural alignment. The highest-ranked alignment corresponds to a basic structural alignment (b) and yields a typical distance error (Sr) of 2.91, a root-mean-square error (Er) of 3.02, 122 structurally equivalent residue pairs (L), and a low sequence identity (IS) of 5%, while allowing for three permutations (red asterisk). The alignment identifies structural equivalence of TMDs forming the hydrophilic back wall of the cavity.

In summary, these results show that although the linear order is shuffled, TMD 1 to 4 of both proteins form a related core structure of the cavity including the hydrophilic backwall. TMD5 of the two proteins do not align and are, because of the inverted membrane topology, appended on different sides of the cavities, resulting in a slightly rotated lateral opening towards the membrane core in the two proteins (Fig. 10A). A very similar structural alignment using TopMatch was obtained for pATOM36 and MTCH2 (Sup. Fig. 5 and 6).

## Discussion

The main finding of this study is that human MTCH1 or N78-MTCH2 can restore the ATOM complex assembly defect in *T. brucei* cells depleted of pATOM36. Previous work demonstrated that *T. brucei* pATOM36 and the yeast MIM complex, composed of Mim1 and Mim2, reciprocally complement the (A)TOM complex assembly defects in their respective systems^30^. More recently, it was reported that MTCH1, but not MTCH2, restores TOM complex assembly in a yeast strain lacking Mim1 and Mim2^16^.

Intriguingly, MTCH2 required the N-terminal extension of MTCH1 to localize to the *T. brucei* mitochondrion and to restore ATOM complex assembly. However, expression of the same MTCH1–MTCH2 fusion protein in a yeast Mim1/Mim2 deletion strain resulted in only partial complementation. This limited rescue may be explained by the observation that, in yeast, the fusion protein, although efficiently inserted into the OMM, was also partially localized to the ER. We also note that the N-terminal extension exhibits a low-complexity, charge-rich composition consistent with an intrinsically disordered region. The functional significance of this feature is currently unknown but may be of interest for future work.

Expression of Mim1 and Mim2^30^, as well as of MTCH1 or MTCH2, only partially restored the growth defect of pATOM36-depleted *T. brucei*. This is due to the dual function of pATOM36, which also serves as an essential subunit of the trypanosomal TAC complex^26^. Because TAC-mediated kDNA segregation is fundamentally distinct from mitochondrial genome segregation in yeast and mammals^28^, this function cannot be complemented by either the MIM complex or by MTCH1 or MTCH2.

Human MTCH2 and the yeast MIM complex insert α-helically anchored proteins into OMM^20,48^. The same is likely the case for pATOM36, even though this has directly been shown for a single substrate only^49^. Thus, we conclude that, despite the lack of sequence homology, the MIM complex, MTCH1/MTCH2, and pATOM36 have the same main function in their respective systems. They insert a subset of α-helically membrane-anchored proteins into the OMM.

Our experimental results together with the AlphaFold3-predicted high quality structures of of MTCH1/MTCH2 and pATOM36 allow for a detailed comparative analysis of the two groups of OMM insertases. A structural comparison with the yeast MIM complex is not possible because it consists of an unknown number of the small Mim1 and Mim2 proteins, each having a single TMD only. Our analysis shows that in line with their analogous function, MTCH1/MTCH2 and pATOM36 share the following features:

1. Both are integral membrane proteins that consist of five TMDs forming a hydrophilic cavity that is open towards the cytosol and sealed on the IMS side.
2. The cavity has an additional lateral opening towards the hydrophobic core of the OMM.
3. The back wall of the cavity is formed by three amphiphilic helices with polar and charged residues towards the cavity, and a U-shaped hydrophobic rim that defines the lateral opening.
4. The five transmembrane domains are arranged in a way that appears to induce local membrane thinning.
5. *In vitro* experiments demonstrate that both insertases have scramblase activity.

The absence of significant sequence similarities and the unique phylogenetic distributions of Mim1/Mim2 (fungi), MTCH1/MTCH2 (vertebrates), and pATOM36 (Kinetoplastids) indicates that the three types of protein insertases are not ancestrally related but arose independently by convergent evolution^20,50^. However, it is possible that even homologous proteins can diverge so much over evolutionary time that their shared ancestry is no longer apparent at the sequence level. In such cases, it can be difficult to distinguish convergent evolution to a similar structure from common ancestry. In the case of MTCH1/MTCH2 and pATOM36, which share a similar structure but no sequence homology, the inverse membrane topology provides an additional strong argument for convergent evolution.

Intriguingly, all five common features mentioned above that are shared between MTCH1/MTCH2 and pATOM36 also extend to YidC, a conserved protein insertase localized in the bacterial cell membrane. YidC mediates membrane insertion of a wide variety of substrates via both Sec-dependent and the Sec-independent pathways and also functions as a chaperone^51–53^. In fact, the structure of YidC shows even more similarities to pATOM36 than it does to MTCH1/MTCH2, because the two proteins share the same topology (Sup. Fig. 7A,B). Unlike, for MTCH1/MTCH2 (Sup. Fig. 7C,D), the C-termini of pATOM36 and YidC are exposed to the same side of the membrane, which corresponds to the side from which the substrates get inserted in OMM or the bacterial cell membrane in their respective systems. YidC is the founding member of the large YidC/Oxa1/Alb3 insertase family found in all three domains of life^54^. It includes YidC and SecY in the (inner) cell membrane of bacteria and archaea. In endosymbiotic organelles, it comprises Oxa1 in the mitochondrial inner membrane and Alb3 in the thylakoid membrane of plastids. Finally, the family also includes the ER insertases Sec61 (an orthologue of prokaryotic SecY)^55^, GET1/2, and EMC^56^ orthologues. While all YidC/Oxa1/Alb3 family members function as lateral insertases for specific subsets of α-helical membrane proteins, SecY and Sec61 also can completely translocate a wide variety of soluble substrates from the cis to the trans side of the membrane. The YidC/Oxa1/Alb3 family not only shares functional, but also structural features, the main one being that they have hydrophilic cavities that, in most cases, are formed by five TMDs^57^. This is not surprising, as they all evolved from a common ancestor. However, it is unexpected that virtually all common features of the independently evolved MTCH1/MTCH2 and pATOM36 insertases listed above are also shared by the YidC/Oxa1/Alb3 family (Sup. Fig. 7). This observation suggests that all of these insertases operate through a similar mechanism. Central to their function is a hydrophilic cavity that opens toward the cis side of the membrane and has a lateral opening facing the hydrophobic interior of the lipid bilayer. Short hydrophilic regions of substrates flanking the TMDs are recruited into this cavity, possibly via electrostatic interactions, enabling them to cross the cis-facing leaflet of the membrane. Subsequently, the substrates are further translocated and inserted into the membrane. These steps depend on a so-called “hydrophobic slide” that lines the lateral opening of the cavity and are facilitated by local membrane thinning induced by the insertases^58^. The presence of this hydrophobic slide may also explain the scramblase activity associated with many, if not all, of these insertases^35^. Notably, the insertases discussed here do not form pores and remain sealed on the trans side of the membrane. SecY and Sec61 are exceptions, as they not only mediate membrane insertion but can also fully translocate soluble proteins, requiring the formation of transient pores to permit complete translocation.

Intriguingly, natural selection to lower the energetic cost of membrane insertion of TMDs has led to the convergence on a broad mechanism across all three insertases families (YidC/Oxa1/Alb3, MTCH1/MTCH2 and pATOM36). Despite their diverse origins, these systems appear to solve the same biophysical problem in a similar way, suggesting that the number of viable solutions is highly limited. The use of a hydrophilic cavity to lower the energetic barrier of membrane insertion may therefore represent the simplest, and possibly the only, mechanism for efficient substrate integration into biological membranes.

A central question in biology concerns the relative roles of contingency and determinism in evolution^59^. The evolutionary biologist Stephen J. Gould argued that evolution is largely driven by contingency. In a famous thought experiment, he proposed that if we could replay “the tape of life,” the organisms that evolve would differ profoundly each time^60^. The evolutionary histories of the YidC/Oxa1/Alb3, MTCH1/MTCH2, and pATOM36 insertases can be viewed as three independent replays of this “tape of life.” Remarkably, each replay resulted in virtually the same solution to the biological problem of inserting TMDs into biological membranes. The three insertase systems therefore provide compelling examples of deterministic evolution at the molecular level across vast phylogenetic distances, challenging Gould’s emphasis on contingency.

The YidC/Oxa1/Alb3 insertase family emerged very early in evolution. Phylogenomic analyses indicate that YidC and SecY were already present in the last universal common ancestor (LUCA), while Sec61 and Oxa1 were established in the last eukaryotic common ancestor (LECA). In contrast, the OMM insertases Mim1/Mim2, MTCH1/MTCH2, and pATOM36 arose much later, within the lineages leading to fungi, kinetoplastids, and mammals, respectively.

Given that all eukaryotes possess mitochondria and require the insertion of α-helically anchored OMM proteins, it is highly likely that every eukaryotic lineage employs a dedicated OMM insertase. We therefore anticipate that additional, independently evolved OMM insertases remain to be discovered. If the general mechanism outlined above indeed represents the simplest, or only, solution for TMD insertion, these insertases are predicted to share common structural and mechanistic features. Notably, even Mim1/Mim2 may conform to this framework, making the determination of MIM complex stoichiometry and structure an important goal for future studies.

## Supporting Information

The article contains supporting information.

## Funding statement

Work in the labs of A.S. was supported in part by NCCR RNA & Disease, a National Centre of Competence in Research (grant number 205601) and by project grant SNF 205200 both funded by Swiss National Science Foundation. Work in the lab of C.v.B. was supported by the Swiss National Science Foundation project grant SNF 212304.

## Conflict of interest

The authors declare that there are no conflicts of interest with the contents of this article.

## CRediT author statement

S.B: Conceptualization, investigation, data analysis, visualization, writing-original draft and editing

A.S.: Conceptualization, data analysis, writing – original draft, review and editing, project administration and funding acquisition

C.v.B: Conceptualization, data analysis, writing – review and editing, project administration and funding acquisition

## Material and Methods

### Transgenic cell lines

Transgenic procyclic cell lines derived from *T. brucei* strains 427 or 29-13^26,61^ were cultured at 27 °C in SDM-79 medium supplemented with 10% (v/v) fetal calf serum (FCS). The pATOM36 (Tb427.7.5700) 3ʹUTR RNAi cell line, originally generated by Käser et al. (2016), was subsequently adopted for use in this study^26^. RNAi was carried out using pLew100-derived stem-loop vectors encoding a blasticidin resistance marker, which were designed to target nucleotides 88–371 downstream of the pATOM36 stop codon. Tetracycline-inducible expression of pATOM36 (Tb.427.7.5700), human MTCH1 (Q9NZJ7), human MTCH2 (Q9Y6C9), and the fusion protein N78-MTCH2 was achieved by cloning the respective full-length open reading frames (ORFs) into a modified pLew100 expression vector conferring puromycin resistance. The terminal stop codon was positioned to allow expression of a C-terminal triple HA tag or, in untagged constructs, to prevent HA tag expression^61,62^.

### Digitonin extraction

Crude mitochondria-enriched fractions were prepared by digitonin extraction to assess mitochondrial localization of the protein of interest^62,63^. Procyclic cells (1×10⁸) were resuspended in SoTe buffer (20 mM Tris-HCl, pH 7.5, 0.6 M sorbitol, 2 mM EDTA) supplemented with 0.015% (w/v) digitonin to induce selective permeabilization. Following centrifugation at 6,800 x g for 10 min at 4 °C, the mitochondria-enriched pellet was separated from the supernatant. Equivalent cell numbers from each fraction were subsequently analyzed by SDS–PAGE and immunoblotting. In some cases, the mitochondria-enriched pellets were further subjected to detergent-based extraction.

### BN-PAGE immunoblotting and quantification

Mitochondria-enriched fractions were solubilized on ice for 30 min in buffer containing 20 mM Tris-HCl (pH 7.4), 50 mM NaCl, 10% glycerol, 0.1 mM EDTA, and 1 mM PMSF, supplemented with 1% (w/v) digitonin. Insoluble material was removed by centrifugation at 16,000 × g for 15 min at 4 °C, and the resulting supernatants were resolved on 4–15% Mini-PROTEAN® TGX™ precast protein gels (Bio-Rad). Gel electrophoresis was performed using a two-buffer system with a single anode buffer (50 mM Bis-Tris, pH 7.0) and two cathode buffers: Cathode A (50 mM tricine, 15 mM Bis-Tris, 0.12 μM Coomassie Brilliant Blue G-250, pH 7.0) for the first half of the gel, and Cathode B (50 mM tricine, 15 mM Bis-Tris, pH 7.0) for the second half. Electrophoresis was carried out at a maximum of 120 V and 15 mA for 3-4 hours. Prior to transfer, gels were equilibrated 5 min in SDS-PAGE running buffer (25 mM Tris, 1 mM EDTA, 190 mM glycine, 0.05% (w/v) SDS) to facilitate protein transfer to PVDF membranes (Thermo Fisher Scientific) in 20 mM Tris, 150 mM glycine, 0.02% SDS, 20% methanol^29,64^. Proteins were detected using an ATOM40-specific primary antibody followed by HRP-conjugated secondary antibody. Chemiluminescent signals were visualized using SuperSignal West Pico Plus or Femto substrates (Thermo Fisher Scientific). Line profile intensities of protein complexes were quantified using the Plot Profile plugin in Fiji (version 1.54p windows), and the corresponding areas under the curve were calculated in GraphPad Prism (version 10.6.1) using the preset trapezoidal integration method.

### Immunofluorescent microscopy

A total of 1×10⁶ exponentially growing cells were collected by centrifugation (2,700 × g, 1 min, room temperature), washed with PBS, and resuspended in 50 μl PBS. Cells were allowed to adhere to glass slides and permeabilized for 30 s in PBS containing 0.2% Triton X-100. Following permeabilization, samples were washed with PBS and fixed in 4% paraformaldehyde for 10 min. Fixed cells were washed and blocked in PBS containing 2% (w/v) bovine serum albumin (BSA) prior to sequential incubation with primary and secondary antibodies diluted in PBS/2% BSA. After antibody labeling, slides were washed with PBS, air-dried, and mounted with Vectashield containing 4ʹ,6-diamidino-2-phenylindole dihydrochloride (DAPI) (Vectorlabs)^29^. The slides were imaged on a Nikon Eclipse Ti2 Super-Resolution Microscope (Nikon) using the preset SD432 nm and SD515 nm confocal fluorescence channels.

### kDNA evaluation

Z-stack images of DAPI stained and fixed cells were acquired using the confocal SD432 nm fluorescence channel. Quantification of kDNA areas was carried out in Fiji (version 1.54p, Windows) using a custom semi-automated macro. In brief, Z-stacks near the focal plane were converted into maximum-intensity projections, followed by background subtraction to improve signal-to-noise ratio. Images were then thresholded using a consistent intensity cutoff to generate binary masks, and kDNA areas were quantified with the “Analyze Particles” function. The kDNA loss was visually assessed by combining brightfield and fluorescent images, including only fully visible, intact cells.

### Heterologous expression and purification of pATOM36

Full-length pATOM36 was cloned into the pDDGFP-Leu2d vector^65^, allowing galactose-inducible expression of a fusion protein bearing a C-terminal TEV protease cleavage site, yeast-enhanced green fluorescent protein (yeGFP), and 8xHis tag. The resulting plasmid was transformed into *Saccharomyces cerevisiae* strain BJ5460 (MATa ura3-52 trp1 lys2-801 leu2Δ1 his3Δ200 pep4::HIS3 prb1Δ1.6R can1 GAL) (ATCC-208285) and plated onto plates lacking uracil. Heterologous expression and detergent-based purification was performed following the protocol described by Drew et al. (2008)^66^. A leucine-deficient medium was used for protein expression (2.0 g l⁻¹ yeast synthetic drop-out medium without Leu (Sigma-Aldrich), 6.7 g l⁻¹ yeast nitrogen base without amino acids (Sigma-Aldrich), and 0.1% glucose). Protein expression and mitochondrial localization were evaluated by fluorescence microscopy, with yeast cells stained with MitoTracker™ Red CMXRos (Thermo Fisher Scientific) at the end of the expression. Confocal images were acquired using the SD515 nm and SD595 nm optical configurations. A detergent mixture consisting of lauryl maltose neopentyl glycol (LMNG) (Anatrace) and cholesteryl hemisuccinate (CHS) (Sigma-Aldrich) at a 10:1 (w/w) ratio was used for protein solubilization and purification.

### Liposome preparation

Lipids comprising 90% POPC (w/w%), 9.5% POPE, and 0.5% NBD-PE (all from Avanti Polar Lipids) were initially solubilized in chloroform, dried under a nitrogen stream, and subsequently subjected to overnight vacuum desiccation^35^. The resulting lipid film was resuspended in measurement buffer (10 mM HEPES pH 7.5, 10 mM NaCl, 10 mM KCl) to generate a stock of 10 mg/ml. To produce unilamellar liposomes, the suspension was subjected to five freeze-thaw cycles using liquid nitrogen and a heat block set at 29.4 °C. The unilamellar liposomes were then extruded 21 times through a 100 nm polycarbonate membrane (Whatman® Nuclepore™) to obtain vesicles of defined size.

### Reconstitution of pATOM36 into liposomes

The liposome solution (10 mg/ml) was initially destabilized with 0.65% n-dodecyl-β-D-maltoside (DDM) (Anatrace), followed by the addition of 0.7 μM purified pATOM36 to generate pATOM36 proteoliposomes (5 proteins/vesicle). For control empty liposomes, an equal volume of measurement buffer (10 mM HEPES pH 7.5, 10 mM NaCl, 10 mM KCl) was added. The reaction was incubated at 4 °C for 30 min. Detergent removal was achieved stepwise by adding 30 mg, 50 mg, and 90 mg (wet weight) of Bio-Beads SM-2 (Bio-Rad) at 30 min intervals at 4 °C on a rotating wheel. The liposome suspension was then subjected to sequential centrifugation, first at 10,000 x g for 10 min, followed by 200,000 x g for 60 min at 4 °C. The resulting liposome pellet was resuspended in 100 µl of measurement buffer.

### Lipid scrambling assay

The *in vitro* lipid scrambling assay was performed essentially as described by Ploier et al. (2016)^67^. For each measurement, 20 µl of liposomes were added to a quartz cuvette (Hellma®) containing 1.5 ml of measurement buffer (10 mM HEPES pH 7.5, 10 mM NaCl, 10 mM KCl). Fluorescence was monitored in real time using a PTI Photon Technology International fluorescence spectrometer with excitation at 470 nm and emission at 530 nm. The reaction was initiated by the addition of 20 mM sodium dithionite and terminated by the addition of 0.5% Triton X-100, which equilibrated the system.

### Antibodies

Antibody dilutions used for immunoblotting (IB) and immunofluorescence (IF) are indicated in parentheses. Polyclonal rabbit pATOM36 (IB, 1:50), ATOM14 (IB, 1:50), ATOM19 (IB, 1:50), ATOM40 (IB, 1:10’000, IF, 1:1’000), ATOM46 (IB, 1:50), and ATOM69 (IB, 1:50) have been previously described^22,23,26^. The commercially available antibodies used in this study were monoclonal mouse anti-HA (Enzo Life Sciences AG, cat. no. CO-MMS-101 R-1000; IB, 1:5’000), polyclonal rabbit anti-MTCH1 (Invitrogen, cat. no. PA5-100201; IB, 1:100), polyclonal rabbit anti-MTCH2 (Proteintech, cat. no. 16888-1-AP, IB, 1:100), and monoclonal mouse anti-EF1α (Merck Millipore, cat. no. 05–235; IB, 1:10’000). Secondary antibodies for IB evaluation were IRDye 680LT goat anti-mouse, IRDye 800CW goat anti-rabbit (both LI-COR Biosciences; IB, 1:10’000), and HRP-coupled goat anti-rabbit (Sigma-Aldrich; IB, 1:4’000). Secondary antibody for IF was goat anti-rabbit Alexa Fluor™ 488 (ThermoFisher Scientific; IF 1:1’000).

### Generation and evaluation of structural protein models

Structural models were predicted using the AlphaFold3 server^38^, oriented computationally within a planar OMM lipid bilayer using the PPM 3.0 web server implemented in the OPM database^39^, and visualized with UCSF ChimeraX (version 1.10.1)^68^. The sequences used for structural modeling comprised pATOM36 (Tb427.7.5700, residues 1–320), MTCH1 (Q9NZJ7, residues 1–389), MTCH2 (Q9Y6C9, residues 1–303), and ATOM19 (Tb927.9.10560, residues 1-168). All tools were applied using default parameters unless stated otherwise. Sequence analyses to identify protein motifs and folds were conducted using the InterPro web tool^69^. Pairwise structural alignments were performed using the TopMatch algorithm (version 7.5)^70^ with default parameters, permitting structural permutations.

## Supplementary Figures

**Supplementary Figure 1:**
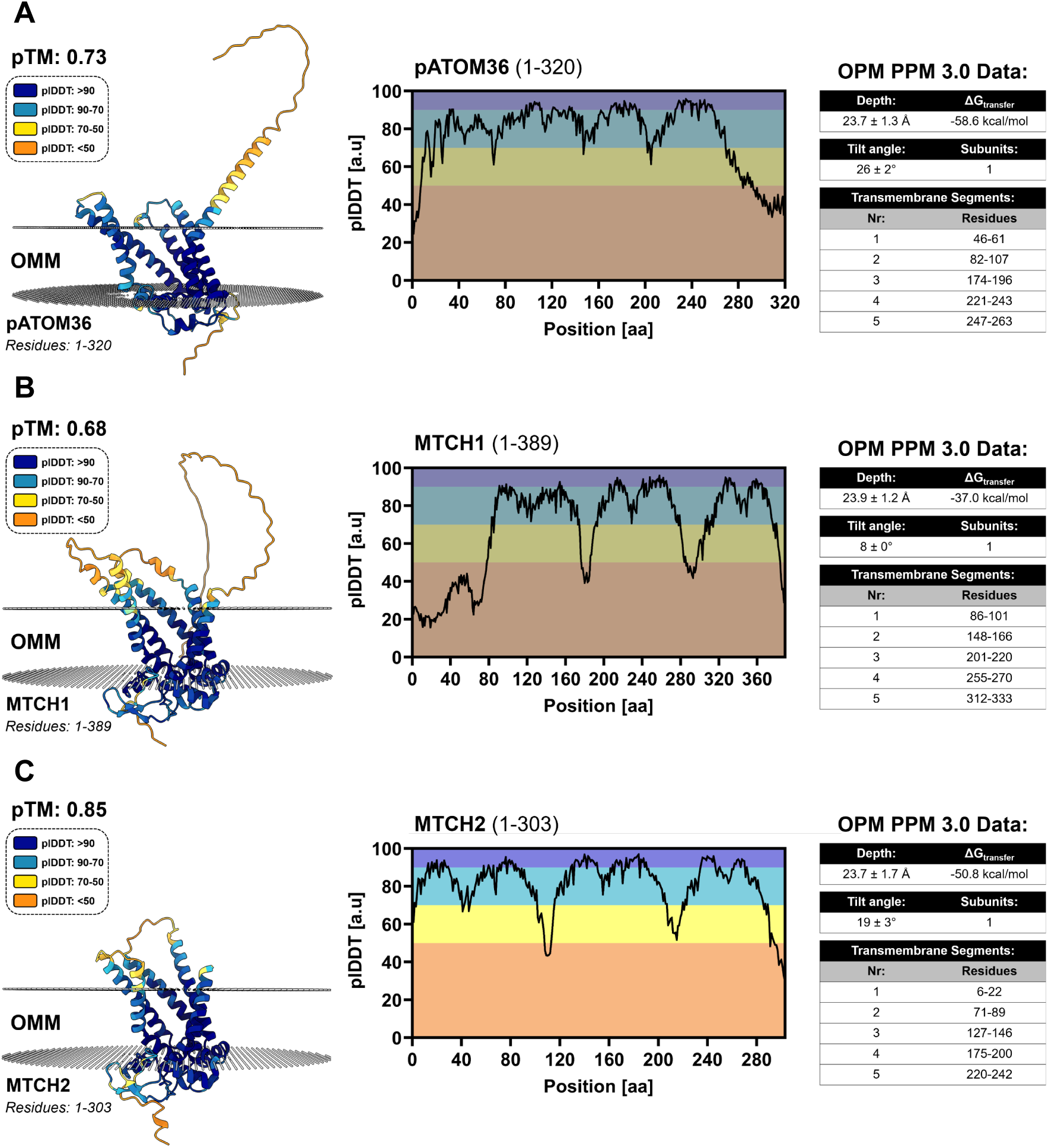
AlphaFold3-predicted structures of OMM protein insertases with confidence assessment and membrane positioning. Structural models were generated using AlphaFold3 and are shown colored by per-residue confidence scores (pLDDT), indicating regions of high and low structural reliability. Residues with pLDDT >90 are shown in dark blue, 90–70 in light blue, 70–50 in yellow, and <50 in orange, and predicted TM-scores (pTM) are indicated. Graphs depict per-residue pLDDT scores across the full-length sequences of the corresponding models. Positioning into the OMM were evaluated using the OPM PPM 3.0 server, providing predictions of membrane depth and hydrophobic thickness, tilt angles, and the free energy of membrane insertion (ΔGtransfer). The complete data set for the **A)** pATOM36 model (Tb427.7.5700), **B)** the MTCH1 model (Q9NZJ7), and **C)** the MTCH2 model (Q9Y6C9).

**Supplementary Figure 2:**
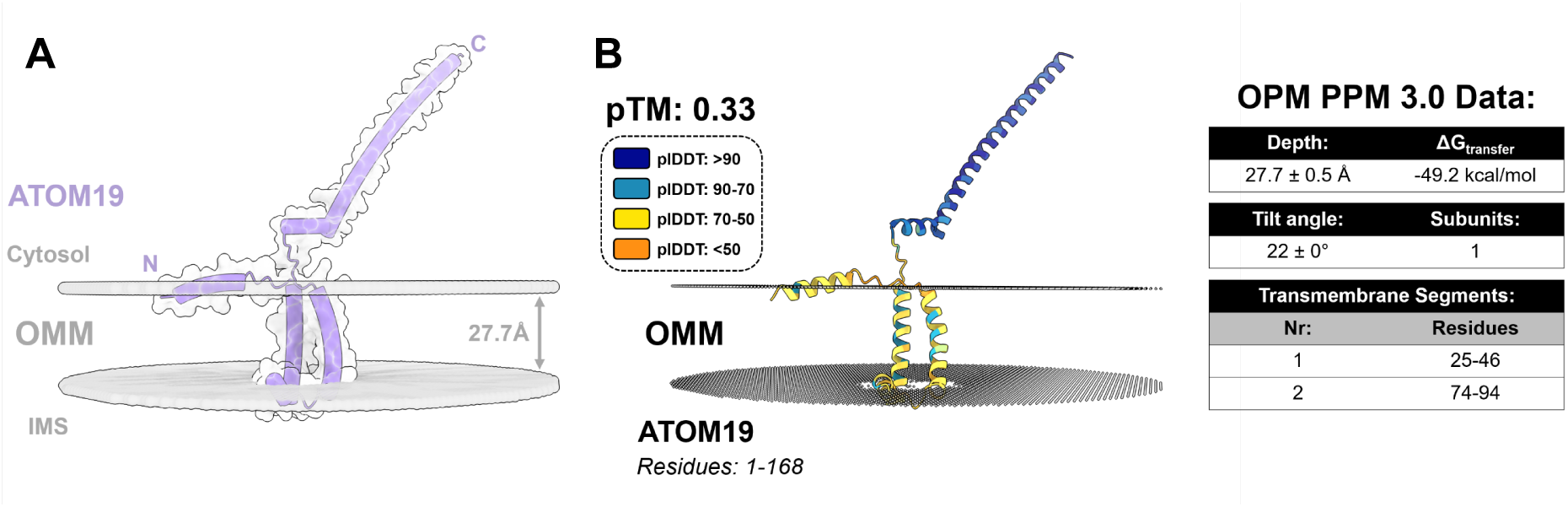
AlphaFold3 model and OMM positioning of the ATOM complex subunit ATOM19. The topology of ATOM19 was predicted using TOPCONS. **A)** Cartoon representation of ATOM19 (lavender) embedded in the OMM (gray), revealing two TMDs and a hydrophobic core thickness of 27.7 Å. **B)** AlphaFold3 confidence-colored model showing per-residue pLDDT scores (>90 dark blue, 90–70 light blue, 70–50 yellow, <50 orange). The pTM value of the entire prediction is indicated. OMM positioning was assessed using the OPM PPM 3.0 server, yielding parameters for membrane depth/hydrophobic thickness, tilt, and free energy of transfer (ΔGtransfer).

**Supplementary Figure 3:**
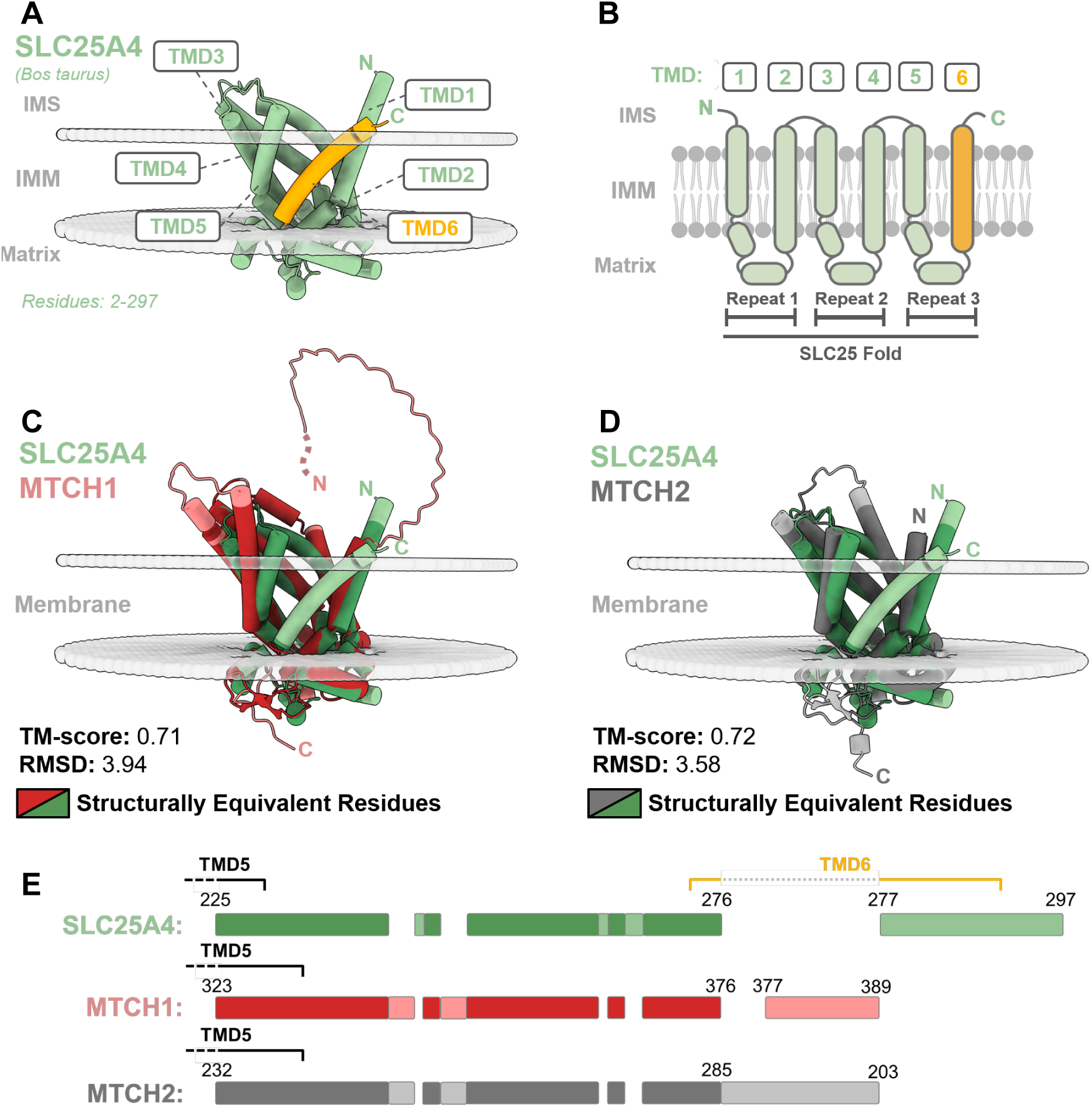
The ADP/ATP carrier (SLC25A4) is a well-characterized member of the SLC25 family. **A)** Crystal structure of the *Bos taurus* SLC25A4 (green) carrier (PDB: 1OKC) at 2.20 Å resolution, embedded in the inner mitochondrial membrane (IMM). The carrier contains six transmembrane domains (TMDs), with TMD6 highlighted in orange. Both the N- and C-termini are oriented toward IMS. **B)** Schematic of SLC25A4 illustrating the canonical SLC25 fold, with six TMDs exhibiting internal threefold pseudo-symmetry. Cartoon representations of the multiple structural alignments between SLC25A4 and **C)** MTCH1 (coral) or **D)** MTCH2 (gray). **E)** Schematic of the jFATCAT rigid (RCS PDB web-tool) alignment, highlighting structurally equivalent residues, SLC25A4 (dark green), MTCH1 (dark red), and MTCH2 (dark gray). Up to TMD5, MTCH1, MTCH2, and SLC25A4 display high structural equivalence. Neither MTCH1 nor MTCH2 shows structural equivalence to TMD6 (orange) of SLC25A4.

**Supplementary Figure 4:**
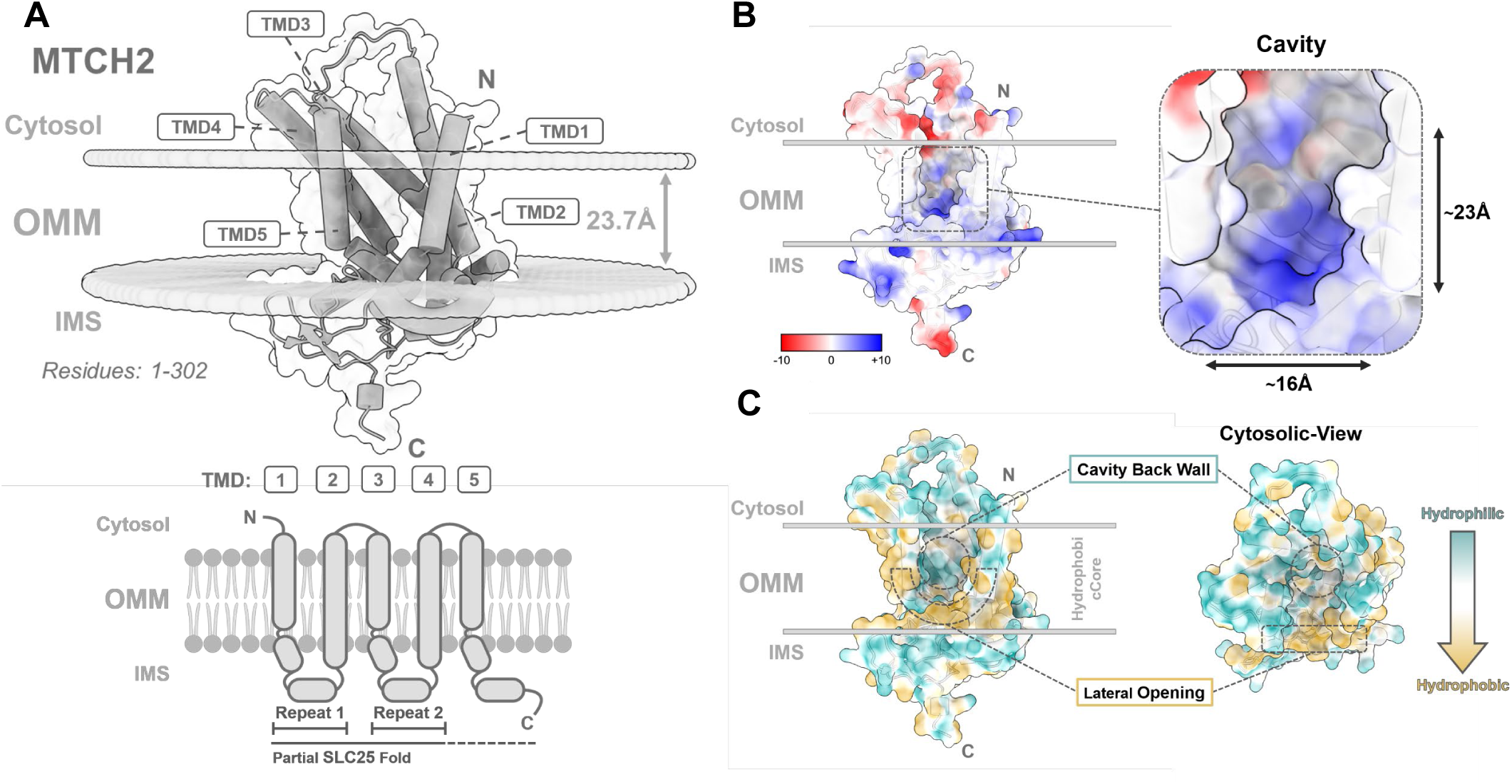
Predicted structural features of human MTCH2. A) AlphaFold3 model of MTCH2 (grey) embedded in the outer mitochondrial membrane (OMM) using the PPM 3.0 server. *Top:* Cartoon representation of residues 1–303, showing five predicted TMDs. The hydrophobic membrane core is reduced to 23.7 Å (grey arrow), with the C-terminus facing the IMS and the N-terminus toward the cytosol. *m:* Schematic of the five TMDs, partially adopting the SLC25 fold. B) Electrostatic surface mapping highlights positively (blue) and negatively (red) charged residues lining a cavity extending ∼23 Å into the hydrophobic membrane core and ∼16 Å wide. **C)** Hydrophobicity e representation shows hydrophobic (brown) and hydrophilic (cyan) regions. Dashed circles indicate the back wall of the cavity and the ped lateral gate. As in pATOM36 and MTCH1, the cytosolic view reveals an amphipathic cavity architecture.

**Supplementary Figure 5:**
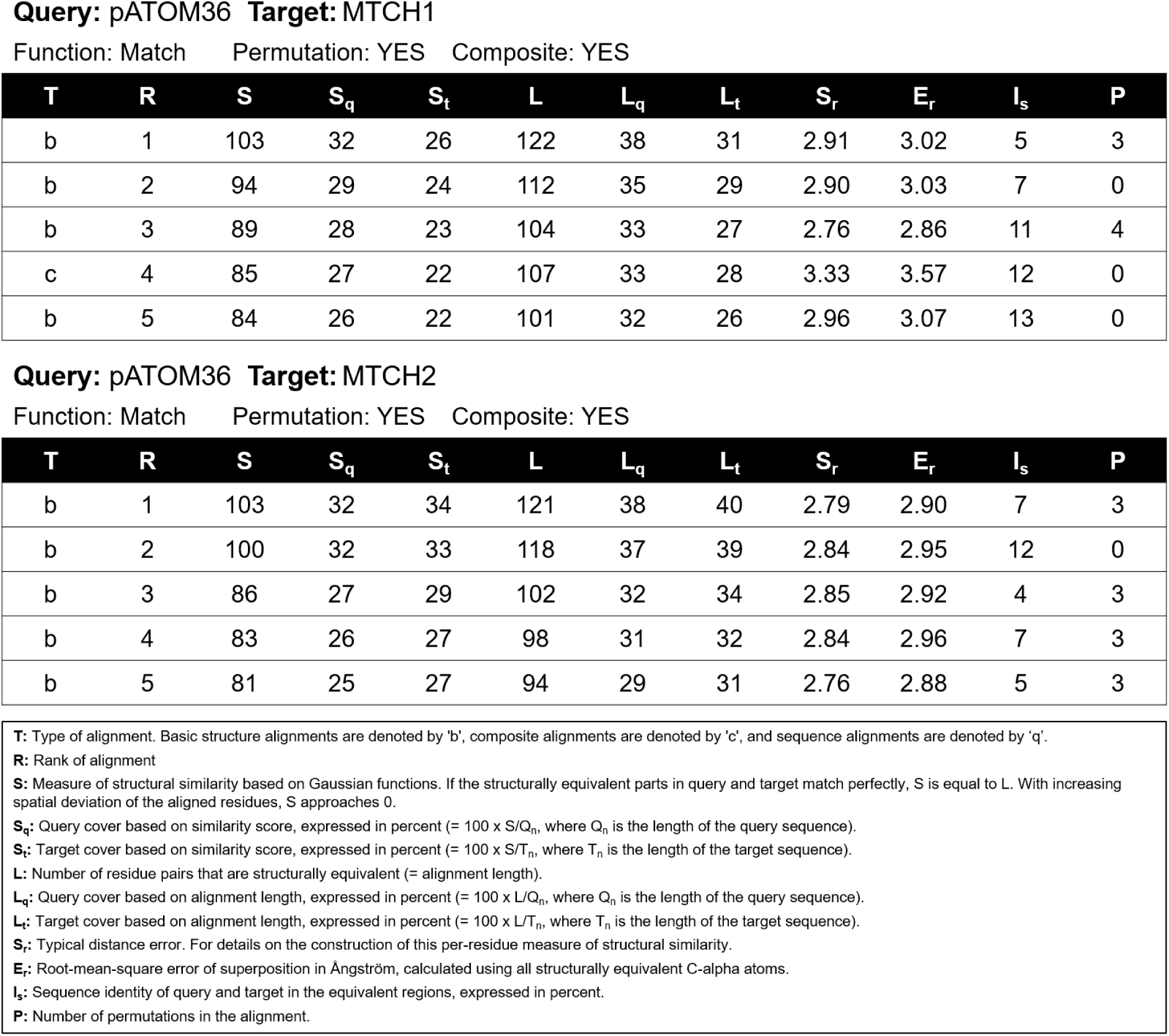
TopMatch pairwise structural alignment of pATOM36 and MTCH1/2. In both analyses, the AlphaFold3 model from pATOM36 was used as the query, while the AlphaFold3 models from MTCH1 and MTCH2 served as the respective targets. Alignments were generated using the match function with permutations and composite type of alignments enabled. The five highest-ranked alignments generated by the algorithm are shown, and the top-ranked structural alignment was selected for each comparison. The individual output parameters contributing to this ranking are defined below.

**Supplementary Figure 6:**
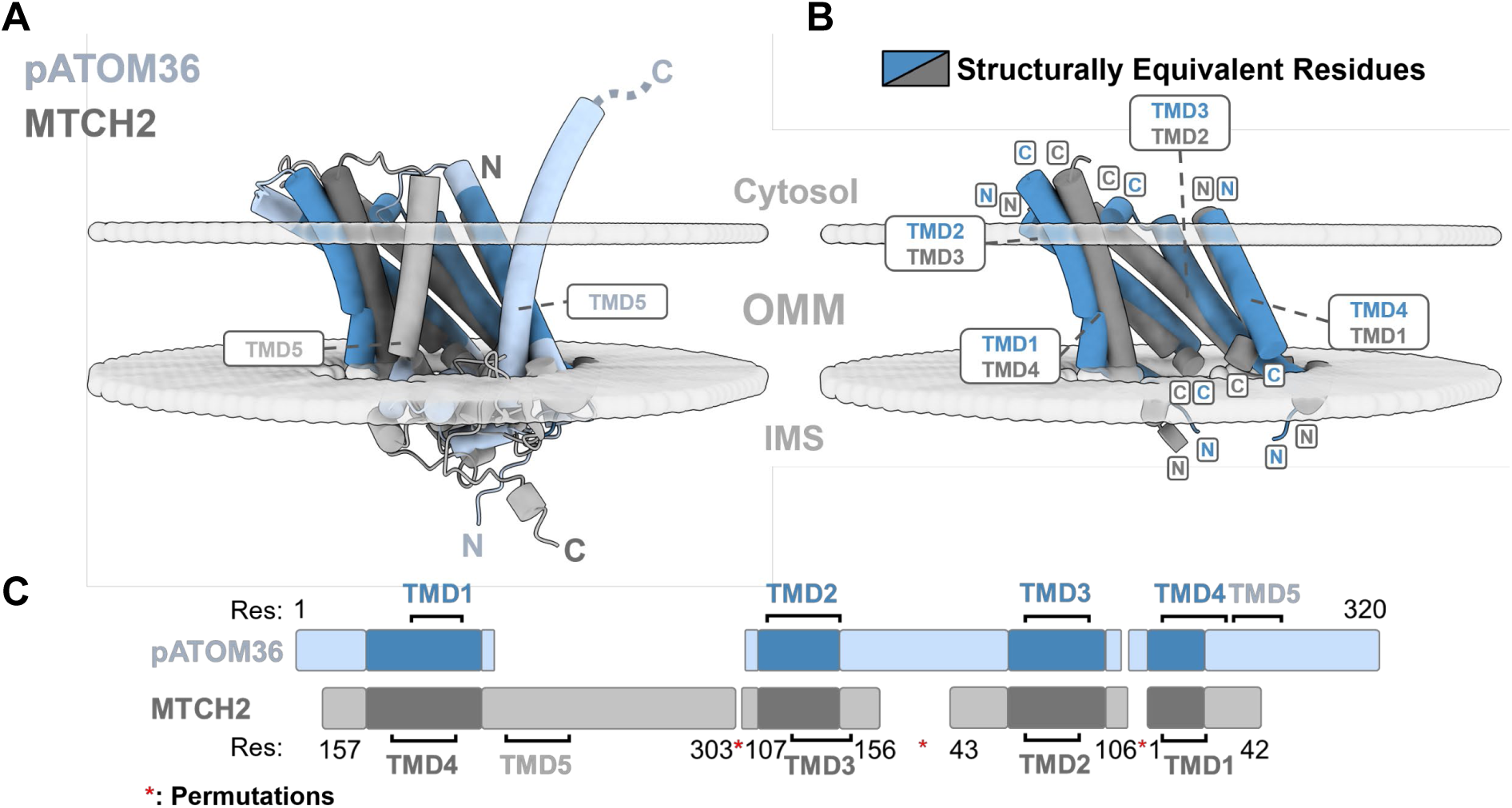
Pairwise structural alignment of pATOM36 and MTCH1. The TopMatch algorithm was used to align pATOM36 (light blue) with MTCH2 (grey). **A)** Global alignment reveals that both proteins adopt inverted topologies while retaining cytosol-accessible cavities. **B)** Structurally equivalent residues between pATOM36 (dark blue) and MTCH2 (dark grey) are primarily located in the TMDs, with each segment preserving its local topology. **C)** Schematic representation of the structural alignment. The highest-ranked basic alignment yields Sr = 2.79, Er = 2.90, L = 121, IS = 7%, and allows three permutations (red asterisk), again revealing structural equivalence of TMDs forming the hydrophilic cavity back wall.

**Supplementary Figure 7:**
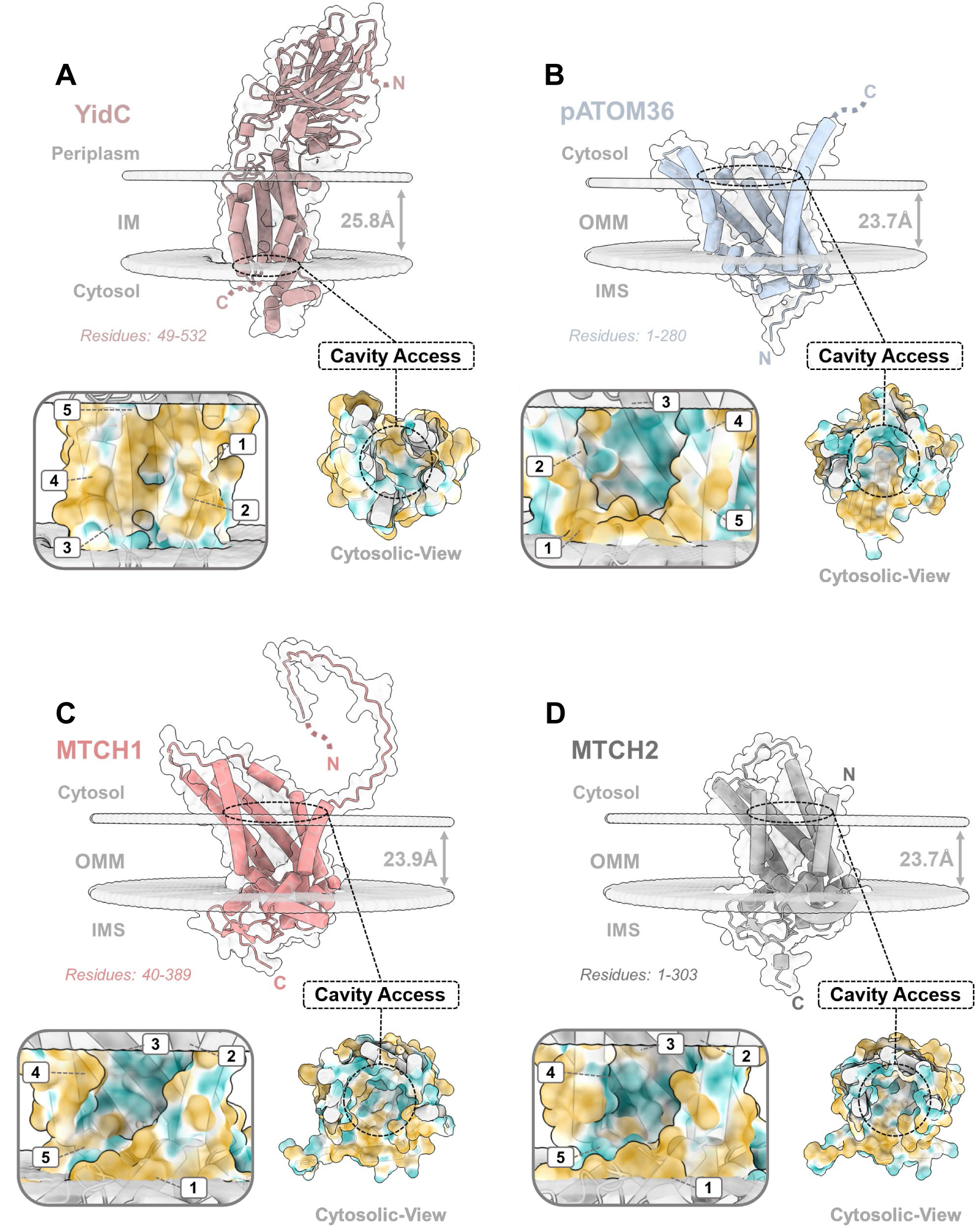
An amphipathic cavity formed by five core transmembrane segments is a conserved structural feature of membrane protein insertases. Structures of representative insertases were analyzed, including the cryo-EM structure of *E. coli* YidC (PDB 6AL2) and AlphaFold3-predicted models of eukaryotic insertases. Hydrophobicity surface representations show that all proteins generate a hydrophilic enivroment within the hydrophobic membrane core. In each case, the amphipathic groove formed by the five transmembrane core segments exhibits a lateral opening toward the lipid phase, remains accessible from the cytosolic side, and is sealed on the posterior side. Shown are **A)** *E. coli* YidC, **B)** *T. brucei* pATOM36, **C)** *H. sapiens* MTCH1, and **D)** *H. sapiens* MTCH2.

